# Language experience predicts music processing in ½ million speakers of 54 languages

**DOI:** 10.1101/2021.10.18.464888

**Authors:** Jingxuan Liu, Courtney B. Hilton, Elika Bergelson, Samuel A. Mehr

**Author notes:** These authors contributed equally (Jingxuan Liu, Courtney B. Hilton).

## Abstract

Tonal languages differ from other languages in their use of pitch (tones) to distinguish words. Lifelong experience speaking and hearing tonal languages has been argued to shape auditory processing in ways that generalize beyond the perception of *linguistic* pitch to the perception of pitch in other domains like music. To examine this, we first conducted a meta-analysis, finding moderate evidence for this idea, but in studies strongly limited by mostly small sample sizes in only a few tonal languages and countries. This makes it challenging to disentangle the effects of linguistic experience from variability in music training experience, cultural differences, and other potential confounds. To address these issues, we used web-based citizen science to test this question on a global scale. We assessed music perception skill in *n* = 34, 034 native speakers of 19 tonal languages (e.g., Mandarin, Yoruba) and compared their performance to *n* = 459, 066 native speakers of other languages, including 6 pitch-accented (e.g., Japanese) and 29 non-tonal languages (e.g., Hungarian). Whether or not participants had taken music lessons, native speakers of all 19 tonal languages had an improved ability to discriminate musical melodies. But this improvement came with a trade-off: relative to speakers of pitch-accented or non-tonal languages, tonal language speakers were also worse at processing the musical beat. These results, which held across tonal languages from a variety of geographic regions and were robust to geographic and demographic variation, demonstrate that linguistic experience shapes music perception ability, with implications for relations between music, language, and culture in the human mind.

## 1 Introduction

From infancy and early childhood, we are surrounded by people speaking and singing (Bergelson et al., 2019; Bonneville-Roussy et al., 2013; Eibl-Eibesfeldt, 1979; Konner, 2010; Mehr et al., 2020, 2019; Mehr, 2014; Mendoza & Fausey, 2021; Yan et al., 2021). This immersion continues throughout the lifespan and is reinforced through our own language and music production.

Human perception readily adapts to these soundscapes: early speech experiences tune our hearing to the speech contrasts of our native language(s) (Kuhl, 2004; Polka & Werker, 1994; Werker & Tees, 1984), and musical experiences during the same time period are thought to have similar “perceptual narrowing” effects, biasing listeners’ interpretations of musical rhythm and pitch based on their own musical cultures (Hannon & Trehub, 2005; Lynch et al., 1990). These effects may cross domains. While music training has minimal causal effects on high-level cognitive skills (Mehr et al., 2013; Sala & Gobet, 2020), it may sharpen lower-level aspects of speech processing (Patel, 2011; Wong et al., 2007) and auditory perception (Kraus & Chandrasekaran, 2010). In the opposite direction, enhanced experience with the types of linguistic pitch used in tonal languages has been argued to shape pitch processing in music (Bidelman et al., 2013; Bradley, 2016; Pfordresher & Brown, 2009).

Here, we study the latter possibility, to examine the effects of language experience on music processing, with a focus on pitch. Languages can be classified into three distinct categories based on their use of pitch: tonal, non-tonal, and pitch-accented. While all spoken languages convey information via pitch, tonal languages, which represent over half the world’s languages (including many East Asian, Southeast Asian and African languages; Yip, 2002) use pitch in a special fashion. In tonal languages, pitch is often used lexically: speaking the same syllable with a different pitch level or shape alters meaning at the word level (Pike, 1948; van der Hulst, 2011). A canonical example is the Mandarin syllable *ma*, which has different meanings depending on its tonal contour (i.e., level, rising, falling-rising, or falling). This property requires pitch sensitivity in both speakers and listeners, lest one scold (*mà*) one’s mother (*mā*) instead of one’s horse (*mă*).

The lexical use of pitch in tonal languages is distinct from how pitch is otherwise used in speech. For example, many languages use pitch to convey affect (Cowen et al., 2019); to cue non-lexical meaning (e.g., helping to differentiate between questions and statements; Patel, 2008; Tong et al., 2005); to emphasize information (Breen et al., 2010); to cue sentence structure with metrical stress patterns (Wagner & McAuliffe, 2019), supporting comprehension (Hilton & Goldwater, 2021); and/or as a cue to speech categories, as in infant- or child-directed speech (Hilton et al., 2022). While these many uses of pitch are typical of speech in both tonal and non-tonal languages (e.g., many Indo-European, South Asian, or Australian languages; Maddieson, 2013), in non-tonal languages, pitch is never used lexically to denote word meanings. Lastly, *pitch-accented* languages form an intermediate category with limited or mixed use of lexical pitch (such as Croatian; van der Hulst, 2011); note, however, that whether pitch-accented languages form a coherent standalone category or whether they are better considered on a spectrum between tonal and non-tonal languages, with some mixed cases, is a matter of debate (e.g., Gussenhoven, 2004; Hyman, 2009, 2006).^1^

The special role of pitch in tonal languages has motivated the hypothesis that speaking a tonal language sharpens pitch perception in a domain-general fashion. Indeed, compared to speakers of non-tonal languages, native speakers of at least some East and/or Southeast Asian tonal languages not only better discriminate the tones of their native language and those of other tonal languages they do not speak (Krishnan et al., 2010; Li & Gao, 2018; Peng et al., 2010), but also may have stronger categorical perception for non-speech pitch patterns generally. Speakers of East/Southeast Asian tonal languages also have distinct neural responses to pitch in brain areas associated with early auditory processing (Bidelman et al., 2011a; Bidelman et al., 2011b; Bidelman & Lee, 2015; Krishnan et al., 2010, 2005).

Might domain-general auditory processing advantages transfer to enhanced pitch processing in music? Many studies have tested this question by comparing native speakers of tonal and non-tonal languages on a variety of musical pitch perception tasks. Some studies report that tonal language speakers excel at discriminating melodic patterns (Alexander et al., 2008; Bradley, 2016; Chen et al., 2016; Choi, 2021; Ngo et al., 2016; Swaminathan et al., 2021; Wong et al., 2012); or at discerning fine-grained pitch difference either in isolation or in the context of detuned intervals, contours, and melodies (Bidelman et al., 2013; Chen et al., 2016; Giuliano et al., 2011; Hutka et al., 2015). But other studies fail to replicate these patterns, both for melodic discrimination (Peretz et al., 2011; Stevens et al., 2013; Zheng & Samuel, 2018) and fine-scale pitch discrimination (Bent et al., 2006; Bidelman et al., 2011a; Pfordresher & Brown, 2009; Stevens et al., 2013; Tong et al., 2018; Wong et al., 2012). Some studies even find that tonal language speakers have *more* trouble distinguishing musical pitch contours, suggesting that lexical tone experience could interfere with pitch perception in some contexts (Bent et al., 2006; Chang et al., 2016; Peretz et al., 2011; Zheng & Samuel, 2018).

More generally, because the vast majority of participants in these studies were native speakers of a small number of tonal and non-tonal languages from two non-overlapping geographic areas (East Asia for tonal languages, with most participants being native speakers of Mandarin or Cantonese; North America for non-tonal languages, with most participants being English speakers), it remains unclear whether patterns of results across these studies reflect effects of tonal vs non-tonal language experience in general; effects of growing up in an East Asian vs Western culture; or some interaction between the two.

In this paper, we first assess the current degree of evidence, via meta-analysis, for an effect of tonal language experience on music processing. Then, we report new data from a massive online experiment that recruited a global sample, to directly measure the relation between linguistic experience and music perception across many tonal, non-tonal, and pitch-accented languages.

## 2 Meta-analytic effects of tonal language experience on music processing ability

We aggregated summary information from 19 prior studies of music perception and tonal language experience (see SI Text 1.1 for the search criteria) and studied them with random-effects meta-analysis models. Because there is no consensus in the literature about what specific music processing ability to expect a tonal-language advantage on, we grouped the prior studies into three rough categories: melody, fine-grained pitch, and rhythm. We built separate meta-analytic models for each group.

The results are in Table 1. The overall effect size estimate suggests that native speakers of tonal languages have overall advantages in both categories of pitch-relevant tasks (melody processing: 0.501, 95% CI = [0.192, 0.81]; fine-grained pitch processing: 0.326, 95% CI = [0.121, 0.53]), but not in rhythm tasks (rhythm processing: −0.008, 95% CI = [-0.14, 0.125]). The meta-analyses also identified three serious concerns, however, which preclude any generalized claim about the effects of tonal language experience on music processing ability.

**Table 1.**
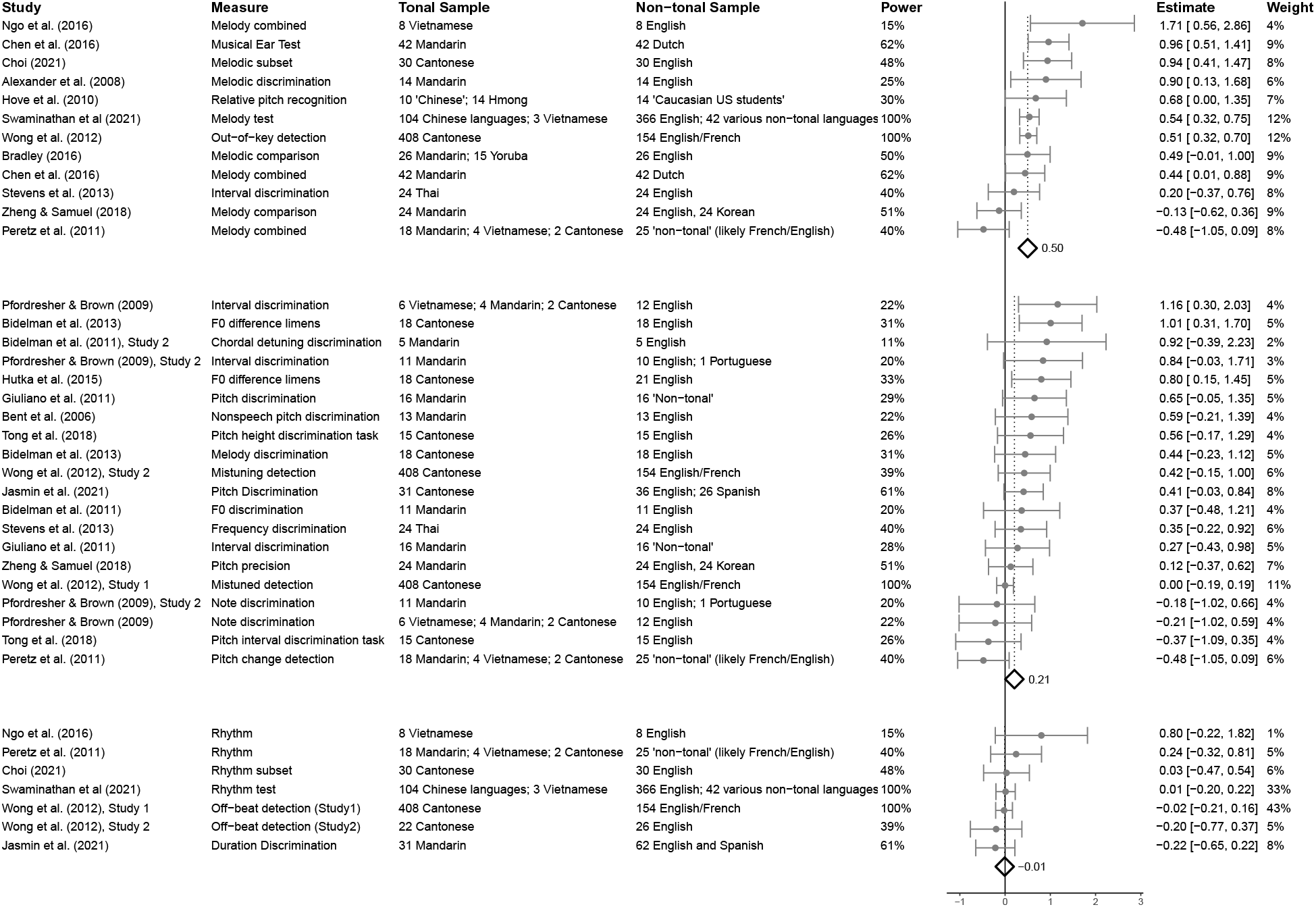
Meta-analytic effects estimated for melody processing, fine-grained pitch processing, and rhythm processing, aggregated from 19 prior studies. Detailed information about the study and measures used, and the language composition of the participants in each study is included in the table. The figure embedded within the table shows the estimated effects for each study individually (modelled as random effects), with 95% confidence intervals indicated by the cross-bars. The bolded diamond shaped points represent the overall fixed-effect estimates for each of the three study categories (melody, fine-grained pitch, and rhythm).

First, and most importantly, prior studies sample tonal languages narrowly, typically comparing Mandarin or Cantonese speakers from mainland China to English speakers from the United States. Of the tonal language speakers in prior studies, approximately 92% spoke Mandarin or Cantonese; and of the non-tonal speakers, approximately 85% spoke English. As such, no claim about the effects of language experience on music perception on the basis of the prior literature is justifiable, because it is not clear whether prior effects generalize beyond a few frequently studied languages.

Second, the majority of prior studies have low statistical power, due their small sample sizes, which may produce unreliable group-level estimates of effects and increase the risk of bias (Kraemer et al., 1998). We estimated the power of each prior study to detect effects of *d* = 0.5, a “medium” size effect comparable to the meta-analytic effect estimated for melodic discrimination tasks (Table 1). Across the three categories of music processing tasks, power was low (for studies of melodic discrimination, median power = 0.49; for fine-grained pitch discrimination, median power = 0.28; for rhythm tasks, median power = 0.48).

Third, participants’ musical training experience has rarely been accounted for in the meta-analysed studies. At best, this contributes additional unsystematic variation within a sample, reducing statistical power. Access to musical training and the form of this musical training, however, may also vary systematically between countries (e.g., Campbell & Wiggins, 2012), potentially leading to biased estimates of music perception abilities. Because the vast majority of participants in prior studies of tonal language experience and music perception ability include native speakers of two languages (i.e., Mandarin and Cantonese) from one country, this issue presents a substantial threat to the validity of prior findings: any systematic biases could produce effects erroneously attributed to differences in tonal language experience rather than cultural experience.

Thus, the meta-analysis demonstrates evidence for a potential effect of tonal language experience on melodic discrimination ability and finer-grained pitch discrimination ability — but a lack of linguistic diversity, low statistical power, and high likelihood of systematic bias in the samples studied warrant caution. These issues can be addressed by studying many native speakers of many languages, with and without music training experience, all of whom complete the same assessments of music processing ability.

In the rest of this paper, we report a study of the ability to discriminate melodies, detect mistuned singing, and detect misaligned beats, in 493,100 people across the globe. Participants self-reported their native language, location, demographics, and degree of musical training, enabling language-wise and language-type-wise analyses of each of these musical abilities and their relations to linguistic and musical experience.

## 3 Citizen-science experiment

Generalization from particular speakers of specific languages to entire groups of languages, with high statistical power, and with controls for potential sources of systematic bias, requires large samples of participants representing a variety of languages and countries (Blasi et al., 2022; Yarkoni, 2022). We report such a test here, using methods of gamification and citizen science (Hartshorne et al., 2019; Hilton & Mehr, 2022; Huber & Gajos, 2020; Long et al., 2023).

### 3.1 Methods

#### 3.1.1 Participants

Participants were visitors to the citizen-science website https://themusiclab.org who completed a set of three music perception tasks presented as an online game (*Test Your Musical IQ*). We did not recruit participants directly; rather, they visited the site after hearing about it organically (e.g., via Reddit posts, YouTube clips, Twitch streams). Participants gave informed consent under an ethics protocol approved either by the Harvard University Committee on the Use of Human Subjects (protocol IRB2017-1206) or the Yale University Human Research Protection Program (protocol 2000033433). Our analysis was preregistered and deviations are noted below. See “Data, code, and materials availability” for more details.

We studied 609,658 participants who completed all three tasks, who had no missing or internally conflicting data, and who reported being a native speaker of one of the 40 most commonly spoken languages among all participants *or* were a native speaker of one of 14 tonal languages that have been less commonly studied in the context of music perception, such as Yoruba, Xhosa, and Lao (n.b., in the preregistration, we initially planned to only study speakers of the 40 most commonly spoken languages in the available data. However, based on advice from reviewers, we subsequently opted to include additional participants to increase the diversity of the tonal languages studied). Data were collected between Nov 8th, 2019 and Nov 12th, 2022.

We excluded participants that (a) had participated in the experiment on another occasion, to avoid any effects of learning (*n* = 33,020); (b) reported a hearing impairment (*n* = 85,963); (c) reported their age as below 8 years or above 90 years (*n* = 1,515); (d) reported an age of music lessons onset that was either below 2 years or above 90 years (*n* = 1,383); (e) reported a music lesson onset age that was greater than their self-reported age (*n* = 370); (f) reported that they were participating in a noisy environment and were not wearing headphones (*n* = 1,910; see SI Text 1.2 for analysis of a manipulation check to test whether participants were actually wearing headphones). 7,107 participants were excluded for meeting more than one of the above criteria. The exclusion criteria were preregistered.

The resulting sample (*n* = 493,100; see Figure 1 and Table S1) included 34,034 native speakers of 19 tonal languages; 16,868 native speakers of 6 pitch-accented languages; and 442,198 native speakers of 29 non-tonal languages.

**Figure 1.**
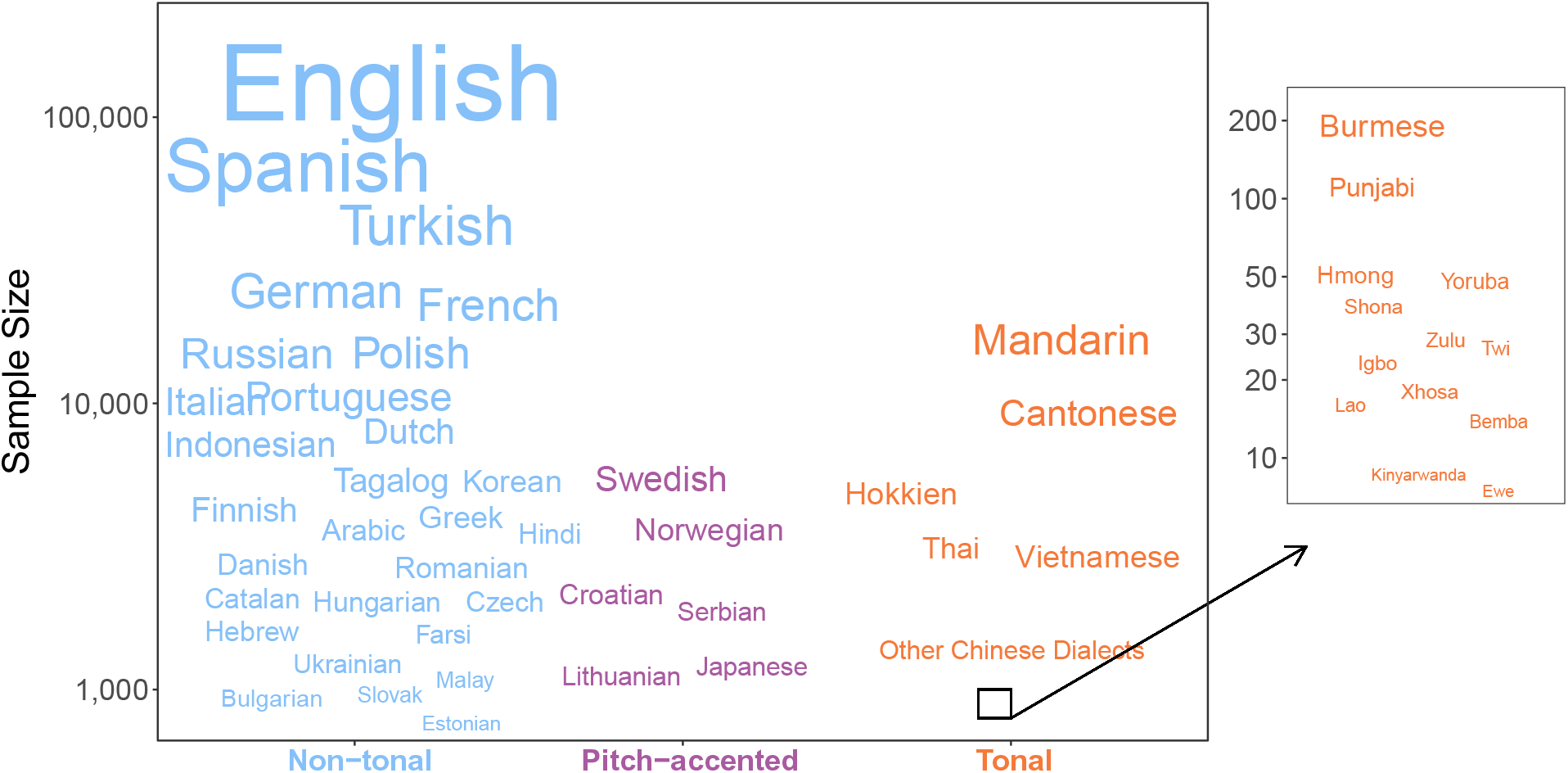
Sample sizes for each language, grouped by language type. The font of each language’s name is scaled proportionally to that language’s sample size. Horizontal positions are jittered to improve readability. The right panel shows additional tonal languages with smaller sample sizes included in the analyses to increase the diversity of the tonal-language sample.

Languages were primarily classified based on the World Atlas of Language Structures (Maddieson, 2013) and the Lyon-Albuquerque Phonological Systems Database (Maddieson et al., 2014). Languages that are not present in either database were classified according to information from the Phonetics Information Base and Lexicon Database (Moran & McCloy, 2019) or other sources from the linguistics literature. A summary of all languages studied here, with further classification details and language-wise sample sizes, is in Table S1.

In addition to participants’ music perception scores, we collected demographic information (gender, age, whether or not the participant had taken music lessons, and the age at onset of those lessons). These data are reported in Table S2.

#### 3.1.2 Stimuli

Participants completed three music perception tasks measuring ability in *melodic discrimination* (Harrison et al., 2017), *mistuning perception* (Larrouy-Maestri et al., 2019), and *beat alignment* (Harrison & Müllensiefen, 2018). The melodic discrimination task assesses the ability to detect differences between melodic patterns: participants listened to three transpositions of the same melody and were asked to choose the version in which one pitch interval was altered (i.e., an oddball task). The mistuning perception task assesses the ability to identify small-pitch differences in vocal pitch: participants listened to two versions of a short musical excerpt, one of which had a vocal track that was detuned from the background music, and were asked to identify the out-of-tune version. The beat alignment task assesses the ability to detect correct synchronization between a click track and some music: participants listened to two versions of the same musical excerpt, both accompanied by a click track; one of the click tracks was misaligned by a constant proportion and participants were asked which example was correctly aligned.

As in the original tasks cited above, each subtask was presented adaptively via `psychTestR` (Harrison, 2020). To minimize the duration of the experiment, we fixed the length of each subtask at 15 trials, the minimum number of trials with acceptably low mean standard errors, according to the original task designs. Demographic items were presented via `jsPsych` (de Leeuw, 2015). The citizen-science platform distributed the experiments using a modified pre-release version of `pushkin` (Hartshorne et al., 2019). Readers can try the three music perception tasks at https://themusiclab.org/quizzes/miq.

#### 3.1.3 Analysis strategy

To test whether musical abilities differ reliably as a function of native language type (i.e., tonal vs. pitch-accented vs. non-tonal), we used mixed-effects linear regression adjusting for age, gender, whether the participant had taken music lessons (yes or no), and the interaction between language-type and music-lessons; with random-effects for language and country. Non-tonal language, female gender, and no-music-lessons were used as the reference levels for the fixed-effects. Stated in terms of {lmer} pseudo-code (Bates et al., 2015), this model is as follows:

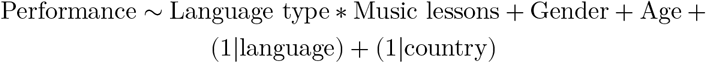

The random-effect structure is particularly helpful for correcting for sampling imbalances, ensuring that no particular languages or countries can dominate the overall effect, while also allowing us to model variation across languages and countries directly (see, e.g., Figure 3).

Note that this analysis approach deviates from our preregistered analysis plan, which involved applying a linear regression model at several sampling levels, without random effects. We made this change on the suggestion of a reviewer and given the utility of mixed-effects models in measuring cultural variation (e.g., Hilton & Mehr, 2022). For transparency, we report analyses and results from the preregistered approach in SI Text 1.2; these largely replicate the results reported in the next section.

### 3.2 Results

#### 3.2.1 Tonal language experience shapes music processing

Native speakers of tonal languages had a reliable advantage in melodic discrimination compared to speakers of non-tonal and pitch-accented languages (Figure 2; full statistical reporting is in Table 2), with an effect size of substantive practical significance (*β*= 0.216, *t* = 4.681, *p* < 0.001), roughly half the size of the effect of having taken music lessons (an experience that one should reasonably expect to directly improve music perception ability). This result replicates the first meta-analytic result, previously shown mainly in Mandarin and Cantonese speakers (see Section 2), demonstrating that the tonal-language advantage for melodic discrimination generalizes to many *additional* tonal languages.

**Table 2.** Fixed-effects from random-effects models for each of the three musical tasks. \*\*\**p* < 0.001, \*\**p* < 0.01, \**p* < 0.05.

| Term | Melodic Discrimination |  |  | Mistuning Perception |  |  | Beat Alignment |  |  |
| --- | --- | --- | --- | --- | --- | --- | --- | --- | --- |
| | $\beta$ | $SE$ | $t$ | $\beta$ | $SE$ | $t$ | $\beta$ | $SE$ | $t$ |
| **Language: Tonal** | **0.216** | **0.046** | **4.681\*\*\*** | **-0.087** | **0.044** | **-1.961** | **-0.225** | **0.032** | **-7.035\*\*\*** |
| Language: Pitch-accented | 0.009 | 0.059 | 0.156 | 0.030 | 0.058 | 0.523 | 0.091 | 0.039 | 2.333* |
| Music lessons: Yes | 0.501 | 0.003 | 147.329*** | 0.417 | 0.003 | 149.319*** | 0.274 | 0.003 | 86.837*** |
| Age | 0.004 | 0.000 | 25.854*** | 0.003 | 0.000 | 21.329*** | -0.002 | 0.000 | -15.374*** |
| Gender: Male | 0.107 | 0.003 | 34.964*** | -0.042 | 0.003 | -16.706*** | 0.142 | 0.003 | 50.006*** |
| Gender: Other | 0.044 | 0.012 | 3.707*** | -0.072 | 0.010 | -7.328*** | 0.046 | 0.011 | 4.095*** |
| Tonal × music lessons | -0.032 | 0.013 | -2.448* | -0.016 | 0.011 | -1.539 | 0.059 | 0.012 | 4.833*** |
| Pitch-accented × music lessons | 0.084 | 0.017 | 5.058*** | 0.055 | 0.014 | 4.009*** | -0.009 | 0.015 | -0.587 |
| Intercept | -0.196 | 0.026 | -7.56*** | 0.113 | 0.025 | 4.526*** | -0.022 | 0.018 | -1.229 |

**Figure 2.**
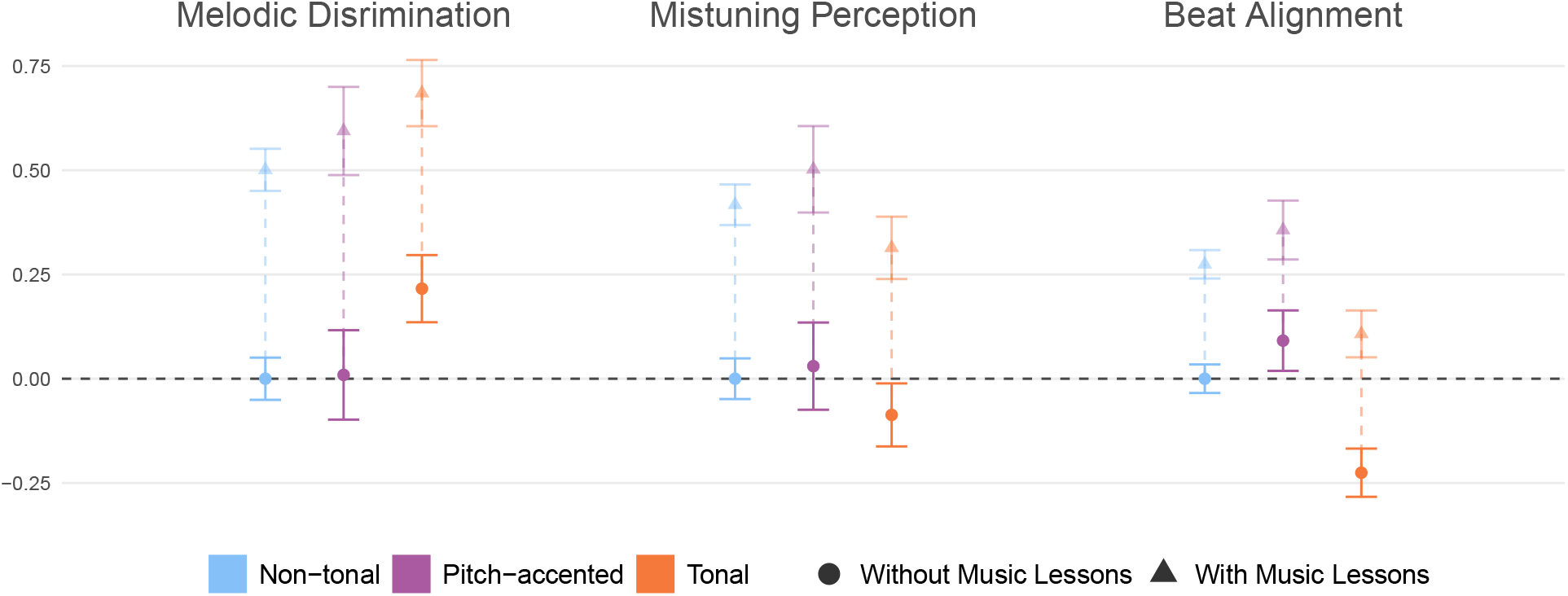
Native speakers of tonal languages have an advantage in melodic discrimination and a disadvantage in beat alignment. The solid dots show the estimated effects of language type, marginalizing over the average proportions for ages and gender, on each of the three music perception abilities tested, but without the estimated effects of having had music lessons. The additive marginal effects of having music lessons are displayed with the faded triangles, providing a comparison point for the language-type effect sizes. After marginalization, for ease of interpretability, a scalar transformation was applied to the coefficients such that the “Non-tonal” and “No music lessons” coefficients were equal to zero. Error-bars represent 95% confidence intervals of the mean. The dotted horizontal black line indicates the baseline (*y* = 0).

Contrasting with the prior meta-analytic effects, however, there was no clear advantage for tonal-language speakers on the vocal mistuning task (*β*= −0.087, *t* = −1.961, *p* = 0.055; Table 2), despite the fact that the task *did* reliably demonstrate clear performance differences between those with and without music training (*β*= 0.417, *t* = 149.319, *p* < 0.001). In other words, while some experiences (e.g., musical training) do shape fine-grained pitch perception in the context of vocal mistuning, there seems to be minimal or no effect of tonal-language experience on this ability. This result contradicts prior studies of fine-grained pitch discrimination that reported tonal-language advantages in mainly Mandarin and Cantonese speakers.

Lastly, contrary the meta-analytic results, which found no effect of tonal language experience on rhythm perception, native speakers of tonal languages performed *worse* in beat perception relative to the non-tonal and pitch-accented groups (*β*= −0.225, *t* = −7.035, *p* < 0.001). Like the melodic discrimination effect, the size of this effect was large, approaching the size of effects of having taken music lessons (see Figure 2).

#### 3.2.2 Language-type effects are consistent across languages

To examine the degree to which the main effects held across the different tonal languages studied, we first examined language-wise estimates derived from the mixed-effects model. These showed a high degree of consistency in main effects within each language group (Figure 3). For example, on the melodic discrimination task, 19 of 19 tonal language groups had an estimated advantage over the non-tonal language average (*ps* < 0.05); whereas 19 of 19 had an estimated disadvantage on the beat perception task (*ps* < 0.05).

**Figure 3.**
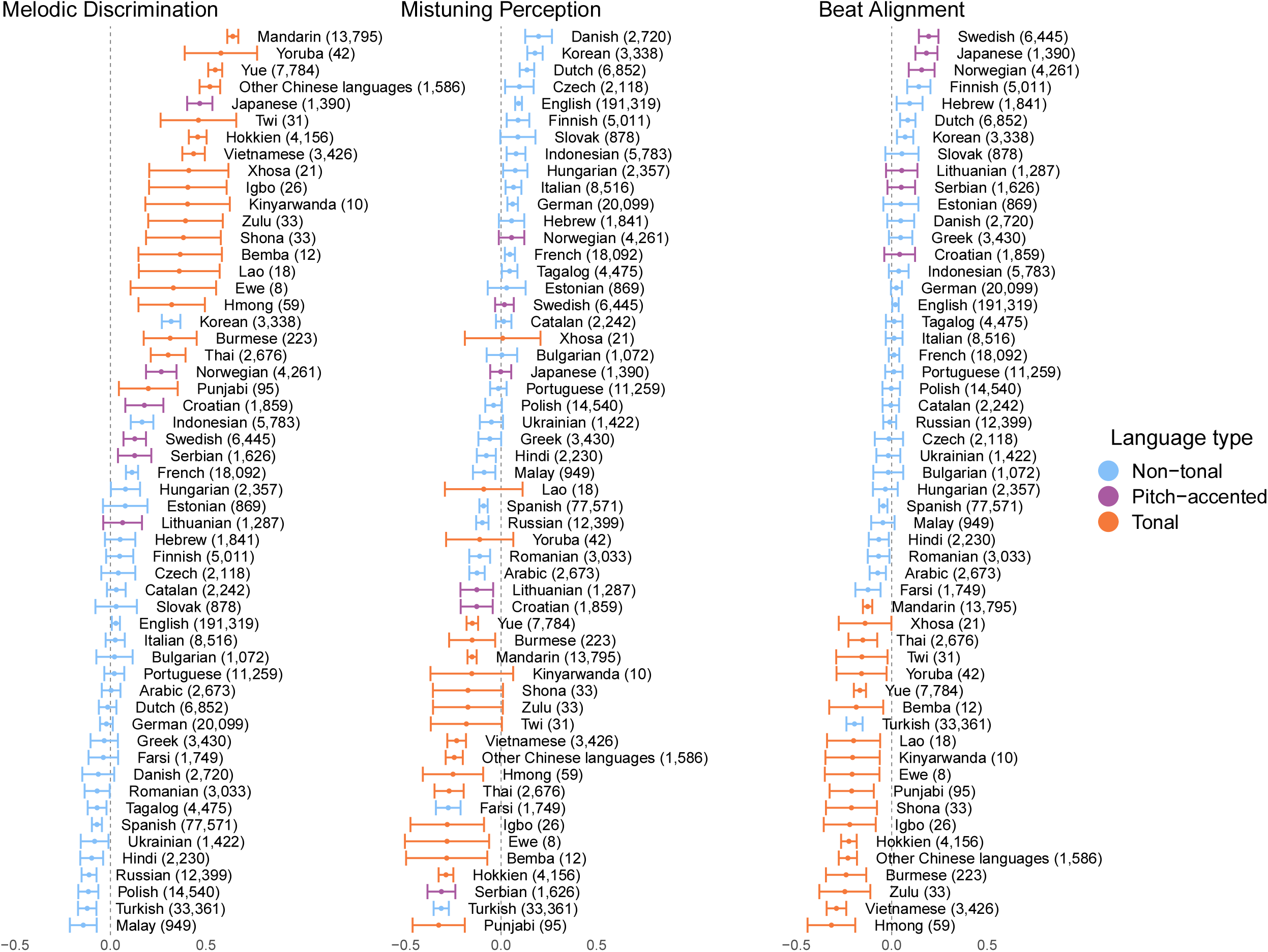
The main effects are consistent across 19 tonal languages. The forest plot displays the estimated average performance for each individual language, after adjusting for the effect of music lessons, age, and gender. The solid points denote random-effect estimates for each language, derived from the mixed-effects models reported in Table 2 and Figure 2; the error bars denote 95% confidence intervals; and the text annotations specify the languages and language-wise sample sizes (in brackets).

We then tested whether speakers of tonal and non-tonal languages could be distinguished on the basis of only their music perception scores, using a linear discriminant function. To compensate for the multi-level structure of our data (scores nested within languages nested within language types), rather than a standard discriminant function analysis, we used a non-parametric permutational approach (Mundry & Sommer, 2007) and ran it on the subset of the participants who indicated having not received musical training and who were native speakers of languages with a sample size of at least 500. From this dataset, we drew nested random samples of 5 tonal languages and 5 non-tonal languages; for each of the sampled languages, we sampled 100 participant scores with replacement and shuffled the assignment of whether that participant was marked “tonal” or “non-tonal”. We trained a linear discriminant function on 30% of that sample, balancing across languages, and then used the trained model to predict whether the remaining 70% of the held-out sample was marked “tonal” or “non-tonal”, yielding a percentage-correct score. We repeated the process 10,000 times to construct the null distribution for each of the three music perception tasks (see Supplementary Figure 2). To estimate the actual classification performance, the same nested sampling process was repeated 100 times for non-shuffled data, from which we obtained the mean proportions of correct classification for each task.

The results robustly replicated the main findings. Speakers of tonal and non-tonal languages could be reliably distinguished on average for both melodic discrimination (*p* < 0.001) and beat alignment (*p* = 0.004) tasks, but not for the mistuning perception task (*p* = 0.099), thus convergently supporting the reliability of our main results.

Last, we used the same linear discriminant analysis approach to compare pitch-accented to non-tonal language speakers (Supplementary Figure 3). Here, the approach failed to replicate the small advantage to pitch-accented language speakers on the beat alignment task (*p* = 0.059) identified by our mixed-effect modeling analysis (Table 2), but did find a small advantage for vocal mistuning (*p* = 0.045). We take this mixed result as cause to not interpret either effect for speakers of pitch-accented languages, given both the less clear theoretical basis for such an effect and the noted ambiguity of the pitch-accented language type (Hyman, 2009).

#### 3.2.3 Effects of tonal-language experience are not attributable to measured third variables

We tested whether the observed language-type effects were driven by systematic variability across participants of three types.

First, we tested whether the socioeconomic status of participants was associated with their task scores, as socioeconomic status may vary systematically with both country and native language in our data, and may mediate the opportunities a person has for musical development. For participants who reported living in the United States, we collected additional demographic information. To assess how socioeconomic status may mediate the main findings, we analysed this subset of participants who reported income information (*n* = 82,727). Despite being restricted to the United States, this sub-sample was nonetheless linguistically diverse, including 3,335 speakers of 14 tonal languages, 189 speakers of 6 pitch-accented languages, and 79,203 speakers of 29 non-tonal languages. We analysed these data using a mixed-effect model of the same structure used in the main analysis, except with an additional fixed-effect term for income. The results show that while income does positively predict performance, such that those with higher incomes tended to perform better, the tonal language melodic discrimination advantage and beat alignment disadvantage held robustly after accounting for these income effects (full statistical reporting is in Table S3).

Second, we examined whether systematic variation in education between speakers of different languages may have driven the observed differences in their performance. Among participants who reported their educational background (*n* = 477,906), we again conducted a mixed-effect analysis of the same structure as the main analysis with an additional fixed term for participants’ education level. While education positively predicted performance on all three tasks, the main language-type effects replicated (Table S4).

Last, we tested whether proximity to “Western” culture could explain the main findings, as the mistuning perception and beat alignment task stimuli used Western-style popular music; while globalized musical styles make this unlikely, it is possible that the degree of familiarity with this musical style systematically varies between tonal and non-tonal language speakers, which could confound the main findings. We compared the results of the main model in two subsets of the participants (combined *n* = 211,256): those who resided in a primarily non-English-speaking Eastern country (China, Taiwan, Hong Kong, Thailand, Vietnam) vs. an English-speaking Western country (United States, United Kingdom, New Zealand, Australia, and Canada). The main language-type effects replicated and the results (Table S5) showed only a small, significant effect of region (West vs. East) on the mistuning perception task.

Taken together, these exploratory analyses show that while SES, education, and exposure to Western culture can correlate with participant performance, these potential confounders cannot account for the observed language-type effects reported here.

## 4 Discussion

We found a clear link between linguistic experience and music processing abilities: native speakers of tonal languages performed better than native speakers of non-tonal languages on a task that required discriminating changes in melodic patterns and worse on a task requiring the perception of a beat. In each case, the effect size associated with being a tonal language speaker was roughly half as large as the effect of receiving music lessons, indicating an effect of substantive practical significance. There was no effect of language experience on the perception of fine-grained pitch, however, despite the fact that musical training was reliably associated with increased ability for this task. This is consistent with the fact that tonal languages do not require distinguishing fine-grained pitch patterns.

Our results are likely to generalize, given that they held across thousands of native speakers of 19 tonal languages, and hundreds of thousands of native speakers of 29 non-tonal languages, each sampled from over 100 countries around the world. The tonal languages include not only East Asian languages (e.g., Mandarin, Cantonese) commonly examined by prior studies, but also Southeast Asian and African languages (e.g., Burmese, Igbo, Shona) that have rarely or never been studied in the context of music perception research. The non-tonal languages in our study include speakers of languages as diverse as Arabic, Catalan, Hindi, and Ukrainian; along with non-tonal languages from South/Southeast Asia, such as Indonesian, Tagalog, and Malay. Inclusion of these Asian languages constitute an especially strong test, since their speakers may be more likely to share cultural similarities to speakers of tonal languages in those regions, thereby helping to rule out cultural confounds.

Our results also help clarify the previously mixed pattern of results concerning the effects of linguistic experience on music processing across different tasks and samples. For example, an advantage for tonal language speakers in melodic pattern processing is consistent with the majority of previous studies (Alexander et al., 2008; Bradley, 2016; Chen et al., 2016; Choi, 2021; Swaminathan et al., 2021; Wong et al., 2012), though not all of them (Peretz et al., 2011; Stevens et al., 2013; Zheng & Samuel, 2018). Meanwhile, similar levels of performance on the fine-grained pitch task between tonal and non-tonal language speakers is supported by some prior studies (Bent et al., 2006; Bidelman et al., 2011a; Stevens et al., 2013; Tong et al., 2018; Zheng & Samuel, 2018) but not other studies that point to a tonal advantage/disadvantage on related tasks (Bidelman et al., 2013; Chang et al., 2016; Giuliano et al., 2011; Hutka et al., 2015; Peretz et al., 2011; Pfordresher & Brown, 2009). Lastly, while rhythmic abilities in tonal language speakers have been less studied (see Wong et al., 2012; Zhang et al., 2020), a disadvantage in beat discrimination is consistent with recent work showing that tonal speakers give more weight to pitch cues than duration cues; this weighting cuts across auditory domains (Jasmin et al., 2021). By leveraging a consistent set of tasks and a large sample size, our results make clear that speaking a tonal language has a measurable connection to music perception ability — but not a uniform one, as this linguistic experience produces both positive and negative effects on perceptual abilities of different types.

Why might tonal language experience have these specific effects on music perception? The answer may lie in the shared mechanisms and neural processing resources associated with auditory perception, whether applied to language or music (Asaridou & McQueen, 2013; Kraus & Chandrasekaran, 2010; Patel, 2008; Patel, 2011; Peretz et al., 2015; although see also Asano et al., 2021; Bidelman et al., 2013). Both tonal languages and music rely on specialized pitch-based sound categories (tone contours or levels in speech; pitch motifs and melodies in music). If these categories are learned and processed through shared, domain-general mechanisms, then improving the efficiency of these mechanisms through practice in either domain should result in mutual improvements (Asaridou & McQueen, 2013 ; Chang et al., 2016; Delogu et al., 2010; Delogu et al., 2006; McMullen & Saffran, 2004; Patel, 2008). But not all is shared. Music relies more on fine-grained pitch structure, even compared to tonal languages, as it is required to support the processing of pitch in the context of a tonal hierarchy (Albouy et al., 2020; Krumhansl, 2004; Peretz et al., 2003; Zatorre et al., 2002). This may explain why tonal language experience did not have an effect on fine-grained pitch processing in music.

Our results do not explain, however, *how* these shared mechanisms might be improved by experience. One possibility is that language experience could shape domain-general perceptual strategies regarding inferences about high-level perceptual categories on the basis of low-level cues: acquired perceptual biases (i.e., from tonal language experience) may aid the processing of some stimuli while worsening the processing of others. In speech, listeners give more perceptual weight to cues that are more informative in discriminating contrasts that are salient in their native language (Jasmin et al., 2021; Schertz & Clare, 2020) and tonal language speakers rely more heavily on pitch to categorize and produce speech stress when acquiring a non-tonal L2 language compared to native speakers (Wang, 2008; Yu & Andruski, 2010).

Similarly, people with pitch perception deficits learn to compensate for their deficits by giving more weight to durational cues when decoding speech prosody (Jasmin, Sun, et al., 2020; Jasmin, Dick, et al., 2020). Recent evidence suggests that similar effects emerge in music perception: Mandarin speakers have difficulty *ignoring* pitch cues relative to English and Spanish speakers, who have been found to more frequently make decisions based on duration cues (Jasmin et al., 2021). In turn, this is consistent with theories of the overlapping mechanisms of basic auditory perception (Patel, 2012, 2011; Peretz et al., 2015; Tierney et al., 2013; Wong et al., 2007). Our findings unite these results and show their generality.

While the scope of our data collection allowed for analysis of music processing abilities in thousands of native speakers of six pitch-accented languages, the findings concerning these speakers were murky and failed to replicate across different analysis approaches. Within-language group variability was also high (see Figure 2) for pitch-accented languages, suggesting no common group advantage/disadvantage across speakers of pitch-accented languages. This, of course, is complicated by the inherently fuzzier nature of classification of pitch-accented languages, relative to tonal languages (see Footnote 1; Gussenhoven, 2004; Hyman, 2009, 2006). Further work that codifies that nature of both linguistic and musical pitch use across this set of generally understudied languages may provide more clarity. We encourage interested readers to re-analyze the open-access data reported here, using alternative classifications of languages into the “pitch-accented” category.

We note several other limitations. First, while we accounted for how much musical training participants had, we did not measure how long they engaged with this training, its intensity, or its type. As a result, our estimates of the effect of musical training have greater uncertainty (although the analyses for participants with no musical training, which largely replicate the main effects, help to mitigate this concern). Second, participants only reported their first language, so we were unable to examine the effects of bilingualism or multilingualism (Krizman et al., 2012; Liu et al., 2020; Liu & Kager, 2017), nor assess whether fluently speaking both tonal and non-tonal languages (e.g. Mandarin and English) might have contributed additional variability in our results. Third, there are a host of other unmeasured cultural, environmental, and genetic factors that surely affect musical abilities. Moreover, these likely interact with each other, complicating causal inferences from the observational data we collected (see, e.g., recent findings that genetics and musical experience both influence linguistic tone perception in Cantonese; Wong et al., 2020). These and other limitations will be best addressed through a variety of methodologies, including more targeted smaller scale approaches (including fieldwork) that complement the broad web-based citizen science approach used here.

In sum, our results show that across a range of geographic and demographic contexts, linguistic experience alters music perception ability in reliable (but not unitary) fashions. This implies that substantively different domains of auditory perception recruit at least some shared processing resources, which themselves are shaped by auditory experience.

## End notes

### Supplemental information

The supplemental information includes 3 text sections, 7 tables and 3 figures.

### Data, code, and materials availability

A reproducible version of this manuscript, including all data and code, is available at https://github.com/themusiclab/language-experience-music. The preregistration is at https://osf.io/xurdb. Readers can try out the experiment at https://themusiclab.org/quizzes/miq; code for each of the three tasks is available at https://github.com/pmcharrison/mpt, https://github.com/pmcharrison/mdt, and https://github.com/pmcharrison/cabat.

## Acknowledgments

This research was supported by the Duke University Internship Funding Program (J.L.); the Harvard Data Science Initiative (S.A.M.); and the National Institutes of Health Director’s Early Independence Award DP5OD024566 (S.A.M. and C.B.H.). We thank the participants; P. Harrison and D. Müllensiefen for sharing code and assisting with the implementation of their music perception tasks; T. Bent, G. Bidelman, E. Bradley, A. Bradlow, D. Chang, W. Choi, R. Giuliano, M. Hove, S. Hutka, M. Ngo, S. Nguyen, I. Peretz, P. Pfordresher, S. X. Tong, S. Swaminathan, Y. Wang, N. Wicha, and L. Zhang for providing supplementary data and/or associated information for the meta-analysis; J. Simson for technical and research assistance; and the members of The Music Lab for discussion and feedback on the citizen-science platform, the experiment, and the manuscript.

## Author contributions

- Conception: J.L. and S.A.M.
- Experimental design and implementation: S.A.M.
- Preregistration and planned analyses: J.L., C.B.H, E.B., and S.A.M.
- Participant recruitment, data management, and data processing: S.A.M., C.B.H., and J.L.
- Analysis and visualization: J.L. and C.B.H., with contributions from E.B. and S.A.M.
- Meta-analysis data collection: J.L.; Meta-analysis models: J.L. and C.B.H.
- Writing: J.L., C.B.H, E.B., and S.A.M.

## Supplementary Information

### 1 Meta-analysis Procedure

#### 1.1 Selection criteria

We included in the meta-analysis only studies that examined the pitch-processing ability of native tonal and non-tonal language speakers via behavioral measures. We searched for studies on Google Scholar using the terms *(tone language OR tonal language) AND (musical pitch perception)* and inspected the first 200 results. In addition, we conducted forward and backward cross-referencing of Asaridou & McQueen (2013), a review article on the link between musical and linguistic pitch. In the identified studies, we excluded those that (1) focused exclusively on absolute pitch, amusia, categorical perception, or cross-modal abilities (e.g. identifying visual representations of musical intervals); (2) studied only musicians; and/or (3) recruited only children under the age of 8. Only studies that were published and written in English were included. In all but one case, the participants studied were native speakers of the tonal language in question. The tonal language group in Swaminathan et al. (2021) contained some non-native speakers; that study’s inclusion or exclusion from the meta-analyses did not substantively affect the estimates of meta-analytic effect sizes, however. Lastly, we excluded one study (Chang et al., 2016) that had a standardized effect size over an order of magnitude above the average effect size. This anomaly was due to suspiciously low standard-deviations of participant-level scores, seemingly due to an accidental shift of a decimal place. We have contacted the authors for clarification.

#### 1.2 Data processing and analysis

To mitigate the confounding effect of musical training, we excluded data from musicians when separate groups of musicians/non-musicians were recruited. The remaining studies that did not distinguish between musicians and non-musicians had participants with either minimal musical training or did not differ across tonal/non-tonal groups ^2^. Additionally, we removed measures regarding pitch memory and speed of processing. For the remaining pitch-relevant tasks, we classified them into two categories: melodic pattern discrimination and fine-tuned pitch discrimination. Melodic discrimination includes tasks that involve recognizing different note combinations, while fine-tuned pitch discrimination includes tasks that concerns discerning fine-grained pitch differences. Additionally, we aggregated rhythm-related tasks (mostly used as control measures) across the studies into a separate rhythm category. Our classification scheme results in 12 melodic discrimination effect sizes, 20 fine-tuned pitch discrimination effect sizes, and 7 rhythm effect sizes. Mean and standard deviation or Cohen’s *d* were collected for all the relevant tasks.

Data from studies/tasks that contained multiple tonal/non-tonal language groups (e.g., separate English and Korean groups) or reported their data at sub-task levels (e.g., 1/4 and 1/2 semitone for the pitch discrimination task) were further processed to produce a composite score for each tonal/non-tonal group or task. Specifically, for studies that contained multiple tonal/non-tonal groups, we used the formula 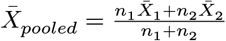 and 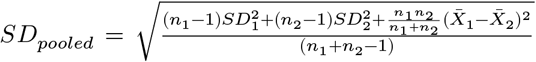 to calculate the combined mean and standard deviation for the group. Meanwhile, for studies that reported data at sub-task levels, we used the formula 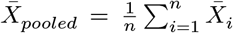 and 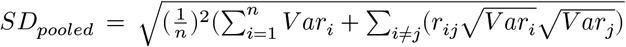 (assuming correlation between the sub-task conditions equals 1, since they are targeting the same ability) to calculate a combined mean and standard deviation for the task for tonal and non-tonal language speakers (following Borenstein et al., 2009). This step produced one entry for each task, which we treat as our unit of analysis.

From the mean and standard deviation, we calculated Hedge’s *G* for each task and used it as our effect size measure. We then ran a random-effects model for each category of the tasks. Please refer to https://github.com/themusiclab/language-experience-music/blob/main/analysis/meta-analysis.Rmd for commented code of the analysis.

### 2 Validation of self-reported headphone use

Participants who self-reported that they were wearing headphones completed a 6-trial headphone detection task (Woods et al., 2017) designed to be easy for participants wearing headphones and difficult for those listening on free-field speakers. Out of the 346,562 participants who indicated wearing headphones, 323,947 had clean and usable headphone detection data. The distribution of scores for these participants (Supplementary Figure 1) was strongly left-skewed with the median participant scoring 5.14 of 6 (100%) correct. This implies that the bulk of participants who self-reported wearing headphones were, in fact, wearing headphones.

### 3 Results from the preregistered analysis approach

In our preregistration (https://osf.io/xurdb), we specified an exploratory-confirmatory approach. For both sets of data, we planned OLS regression models exploratory (*n* = 183,530) and confirmatory (*n* = 307,419) samples, controlling for age, gender, and music lesson (yes or no). In addition, to further reduce the confounding effect of covariates, we planned OLS regression models on three alternative samples, using different approaches to control for differences in music lesson experience, gender, and age (coarsened into 10 year bands). The three versions of the data were a 1:1 exactly matched sample, a 1:1 exactly matched sample with only participants who did not receive music lessons, and an inverse-probability weighted sample (Austin & Stuart, 2015; Stuart, 2010). The same simple linear regression model *Performance* ~ *Language type* was planned for each.Results from the exploratory and confirmatory analyses are presented in Tables S6A & S6B, for transparency. The main findings from the exploratory dataset replicated in the confirmatory dataset.

There are several limitations, however, that complicate the OLS results. First, the large sample size drove almost all effects to statistical significance in a way that may not translate to practical significance. Second, the imbalanced representation of different languages (e.g., dominance of English speakers in the non-tonal language group) may bias our estimates via cultural confounds endemic to the dominant languages in each group. These limitations motivated our adoption of mixed-effect models as the main analysis.

---

**Table S1.** The languages studied here, with sample sizes, largest-sample-size country, and the source of the language classification. Abbreviations: WALS (The world atlas of language structures), LAPSyD (Lyon-Albuquerque phonological systems database).^*^WALS classified as simple tone.

| Language | Language Type | Total *n* | Top country/region | Source |
| --- | --- | --- | --- | --- |
| Chinese/Mandarin | Tonal | 13795 | China | LAPSyD, WALs |
| Chinese/Cantonese | Tonal | 7784 | Hong Kong | WALS |
| Chinese/Hokkien | Tonal | 4156 | Taiwan | WALS |
| Vietnamese | Tonal | 3426 | Vietnam | LAPSyD, WALs |
| Thai | Tonal | 2676 | Thailand | LAPSyD, WALs |
| Chinese/Other dialects | Tonal | 1586 | China | WALS |
| Burmese | Tonal | 223 | Myanmar | WALS |
| Punjabi | Tonal | 95 | India | Evans et al. (2018) |
| Hmong | Tonal | 59 | United States | WALS |
| Yoruba | Tonal | 42 | Nigeria | WALS |
| Shona | Tonal | 33 | Zimbabwe | Jefferies (1990) |
| Zulu | Tonal | 33 | South Africa | Niesler et al. (2009), WALs* |
| Twi | Tonal | 31 | Ghana | Manyah 2006 |
| Igbo | Tonal | 26 | Nigeria | Eme & Odinye (2008)), WALs* |
| Xhosa | Tonal | 21 | South Africa | Niesler et al. (2009) |
| Lao | Tonal | 18 | Laos | Westermeyer & Westermeyer (1977) |
| Bemba | Tonal | 12 | Botswana, Brazil | Hamann & Kula (2015) |
| Kinyarwanda | Tonal | 10 | Rwanda, United States | Goldsmith & Mpiranya (2010) |
| Ewe | Tonal | 8 | Ghana | Ameka (2001), WALs* |
| Swedish | Pitch-accented | 6445 | Sweden | LAPSyD, Bailey (1988) |
| Norwegian | Pitch-accented | 4261 | Norway | LAPSyD, WALs |
| Croatian | Pitch-accented | 1859 | Croatia | Inkelas & Zec (1988); Van der Hulst et al. (2010) |
| Serbian | Pitch-accented | 1626 | Serbia | Inkelas & Zec (1988); Van der Hulst et al. (2010) |
| Japanese | Pitch-accented | 1390 | Japan | LAPSyD, WALs |
| Lithuanian | Pitch-accented | 1287 | Lithuania | LAPSyD |
| English | Non-tonal | 191319 | United States | LAPSyD, WALs |
| Spanish | Non-tonal | 77571 | Spain | LAPSyD, WALs |
| Turkish | Non-tonal | 33361 | Turkey | LAPSyD, WALs |
| German | Non-tonal | 20099 | Germany | LAPSyD, WALs |
| French | Non-tonal | 18092 | France | LAPSyD, WALs |
| Polish | Non-tonal | 14540 | Poland | LAPSyD, WALs |
| Russian | Non-tonal | 12399 | Russia | LAPSyD, WALs |
| Portuguese | Non-tonal | 11259 | Brazil | PHOIBLE, Van der Hulst et al. (2010) |
| Italian | Non-tonal | 8516 | Italy | LAPSyD |
| Dutch | Non-tonal | 6852 | The Netherlands | LAPSyD |
| Indonesian | Non-tonal | 5783 | Indonesia | LAPSyD, WALs |
| Finnish | Non-tonal | 5011 | Finland | LAPSyD, WALs |
| Tagalog | Non-tonal | 4475 | Philippines | LAPSyD, WALs |
| Greek | Non-tonal | 3430 | Greece | LAPSyD, WALs |
| Korean | Non-tonal | 3338 | South Korea | LAPSyD, WALs |
| Romanian | Non-tonal | 3033 | Romania | WALS |
| Danish | Non-tonal | 2720 | Denmark | PHOIBLE, Van der Hulst et al. (2010) |
| Arabic | Non-tonal | 2673 | Egypt | LAPSyD, WALs |
| Hungarian | Non-tonal | 2357 | Hungary | LAPSyD, WALs |
| Catalan | Non-tonal | 2242 | Spain | LAPSyD, WALs |
| Hindi | Non-tonal | 2230 | India | LAPSyD, WALs |
| Czech | Non-tonal | 2118 | Czech Republic | LAPSyD |
| Hebrew | Non-tonal | 1841 | Israel | LAPSyD, WALs |
| Farsi | Non-tonal | 1749 | Iran | LAPSyD, WALs |
| Ukrainian | Non-tonal | 1422 | Ukraine | PHOIBLE, Van der Hulst et al. (2010) |
| Bulgarian | Non-tonal | 1072 | Bulgaria | LAPSyD, WALs |
| Malay | Non-tonal | 949 | Malaysia | PHOIBLE |
| Slovak | Non-tonal | 878 | Slovakia | PHOIBLE, Van der Hulst et al. (2010) |
| Estonian | Non-tonal | 869 | Estonia | Van der Hulst et al. (2010) |

**Table S2.**
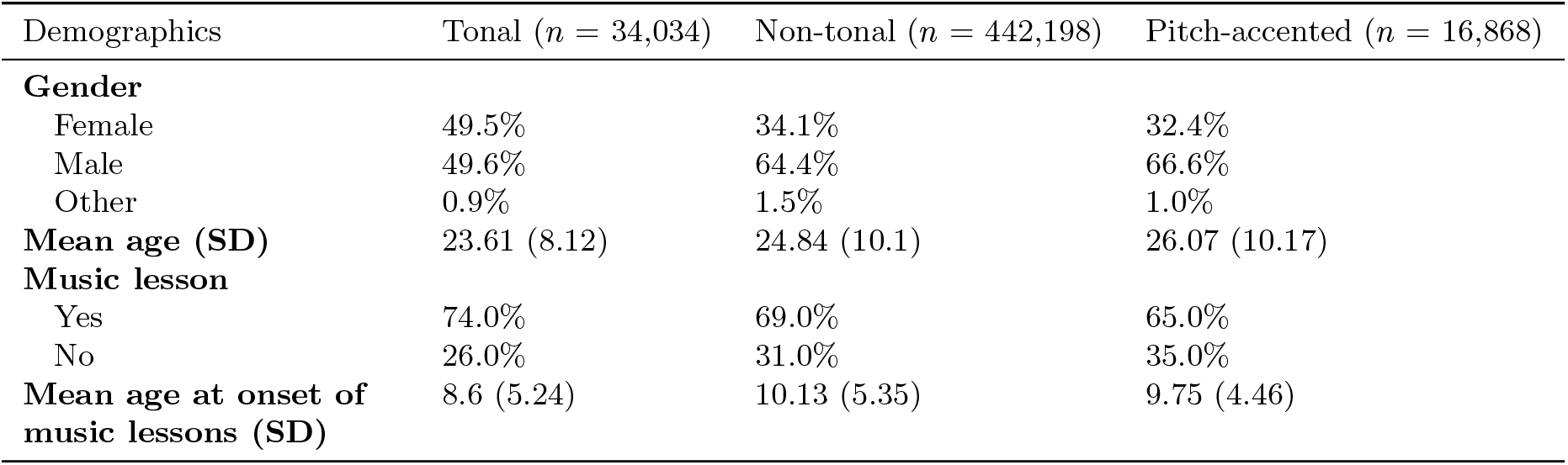
Demographics composition, by language type.

| Demographics | Tonal ($n = 34,034$) | Non-tonal ($n = 442,198$) | Pitch-accented ($n = 16,868$) |
| --- | --- | --- | --- |
| **Gender** |  |  |  |
| Female | 49.5% | 34.1% | 32.4% |
| Male | 49.6% | 64.4% | 66.6% |
| Other | 0.9% | 1.5% | 1.0% |
| **Mean age (SD)** | 23.61 (8.12) | 24.84 (10.1) | 26.07 (10.17) |
| **Music lesson** |  |  |  |
| Yes | 74.0% | 69.0% | 65.0% |
| No | 26.0% | 31.0% | 35.0% |
| **Mean age at onset of music lessons (SD)** | 8.6 (5.24) | 10.13 (5.35) | 9.75 (4.46) |

**Table S3.** Fixed-effect output from random-effects model applied to each musical task controlling for participant annual income (Baseline = Income under 10,000) in the US sample with available income data. Language type effect remains after controlling for income in this group. Language is included as random-effects. \*\*\**p* < 0.001, \*\**p* < 0.01, *p < 0.05.

| Term | Melodic Discrimination |  |  | Mistuning Perception |  |  | Beat Alignment |  |  |
| --- | --- | --- | --- | --- | --- | --- | --- | --- | --- |
| | $\beta$ | $SE$ | $t$ | $\beta$ | $SE$ | $t$ | $\beta$ | $SE$ | $t$ |
| **Language: Tonal** | **0.310** | **0.074** | **4.189\*\*\*** | **-0.162** | **0.068** | **-2.372\*** | **-0.233** | **0.047** | **-4.927\*\*\*** |
| Language: Pitch-accented | -0.024 | 0.179 | -0.132 | -0.152 | 0.145 | -1.045 | 0.219 | 0.146 | 1.499 |
| Music lessons: Yes | 0.577 | 0.009 | 62.859*** | 0.441 | 0.007 | 62.793*** | 0.278 | 0.008 | 34.94*** |
| Age | 0.001 | 0.000 | 3.634*** | 0.001 | 0.000 | 2.027* | -0.004 | 0.000 | -14.458*** |
| Gender: Male | 0.109 | 0.007 | 14.851*** | -0.046 | 0.006 | -8.198*** | 0.118 | 0.006 | 18.538*** |
| Gender: Other | 0.066 | 0.027 | 2.466* | -0.043 | 0.020 | -2.089* | 0.102 | 0.023 | 4.396*** |
| Income: \$10,000 to \$19,999 | 0.037 | 0.019 | 1.899 | 0.022 | 0.015 | 1.472 | 0.036 | 0.017 | 2.138* |
| Income: \$20,000 to \$29,999 | 0.024 | 0.018 | 1.329 | 0.055 | 0.014 | 3.894*** | 0.076 | 0.016 | 4.819*** |
| Income: \$30,000 to \$39,999 | 0.030 | 0.018 | 1.662 | 0.032 | 0.014 | 2.335* | 0.079 | 0.016 | 5.039*** |
| Income: \$40,000 to \$49,999 | 0.044 | 0.018 | 2.4* | 0.075 | 0.014 | 5.421*** | 0.088 | 0.016 | 5.589*** |
| Income: \$50,000 to \$74,999 | 0.045 | 0.016 | 2.852** | 0.070 | 0.012 | 5.838*** | 0.082 | 0.014 | 6.048*** |
| Income: \$75,000 to \$99,999 | 0.062 | 0.016 | 3.859*** | 0.086 | 0.012 | 6.956*** | 0.078 | 0.014 | 5.6*** |
| Income: \$100,000 to \$150,000 | 0.093 | 0.015 | 6.144*** | 0.103 | 0.012 | 8.865*** | 0.096 | 0.013 | 7.323*** |
| Income: Over \$150,000 | 0.121 | 0.015 | 7.897*** | 0.087 | 0.012 | 7.385*** | 0.087 | 0.013 | 6.54*** |
| Pitch-accented × music lessons | 0.207 | 0.188 | 1.105 | 0.217 | 0.145 | 1.498 | 0.016 | 0.160 | 0.099 |
| Tonal × music lessons | -0.044 | 0.043 | -1.028 | 0.047 | 0.033 | 1.411 | 0.089 | 0.037 | 2.396* |
| Intercept | -0.150 | 0.040 | -3.779*** | 0.132 | 0.037 | 3.603*** | 0.032 | 0.026 | 1.263 |

**Table S4.**
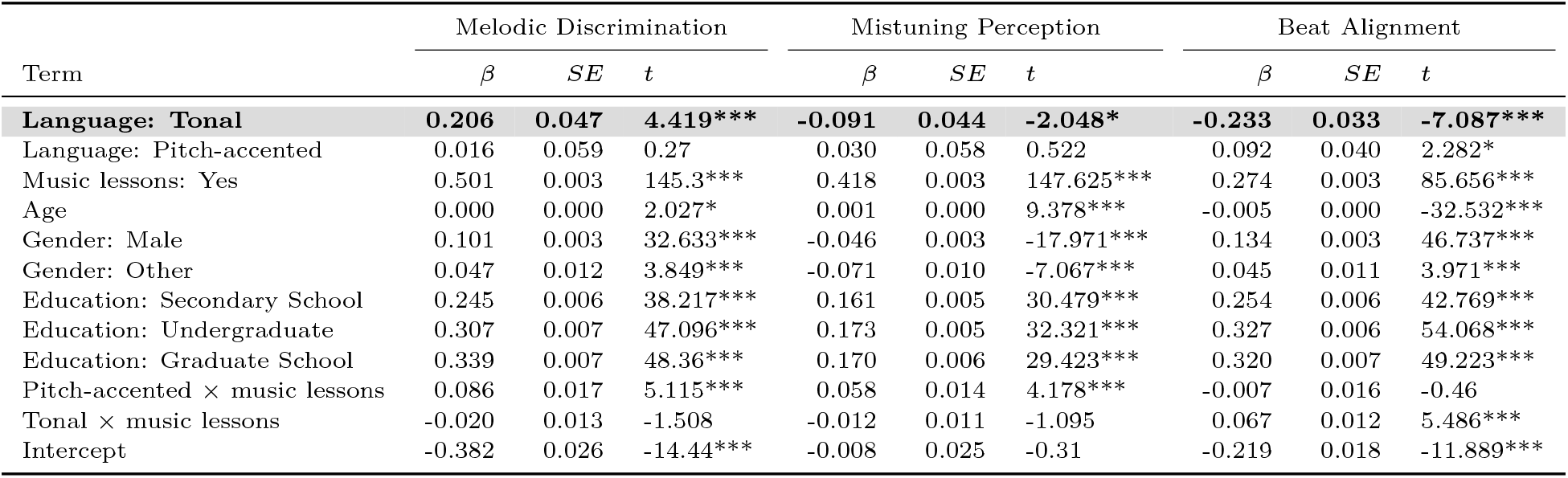
Fixed-effect output from random-effects model applied to each musical task controlling for education (Baseline = Primary School) in the sample that reported their educational background. Language and country are included as random-effects. \*\*\**p* < 0.001, \*\**p* < 0.01, *p < 0.05.

| Term | Melodic Discrimination |  |  | Mistuning Perception |  |  | Beat Alignment |  |  |
| --- | --- | --- | --- | --- | --- | --- | --- | --- | --- |
| | $\beta$ | $SE$ | $t$ | $\beta$ | $SE$ | $t$ | $\beta$ | $SE$ | $t$ |
| **Language: Tonal** | **0.206** | **0.047** | **4.419\*\*\*** | **-0.091** | **0.044** | **-2.048\*** | **-0.233** | **0.033** | **-7.087\*\*\*** |
| Language: Pitch-accented | 0.016 | 0.059 | 0.27 | 0.030 | 0.058 | 0.522 | 0.092 | 0.040 | 2.282* |
| Music lessons: Yes | 0.501 | 0.003 | 145.3*** | 0.418 | 0.003 | 147.625*** | 0.274 | 0.003 | 85.656*** |
| Age | 0.000 | 0.000 | 2.027* | 0.001 | 0.000 | 9.378*** | -0.005 | 0.000 | -32.532*** |
| Gender: Male | 0.101 | 0.003 | 32.633*** | -0.046 | 0.003 | -17.971*** | 0.134 | 0.003 | 46.737*** |
| Gender: Other | 0.047 | 0.012 | 3.849*** | -0.071 | 0.010 | -7.067*** | 0.045 | 0.011 | 3.971*** |
| Education: Secondary School | 0.245 | 0.006 | 38.217*** | 0.161 | 0.005 | 30.479*** | 0.254 | 0.006 | 42.769*** |
| Education: Undergraduate | 0.307 | 0.007 | 47.096*** | 0.173 | 0.005 | 32.321*** | 0.327 | 0.006 | 54.068*** |
| Education: Graduate School | 0.339 | 0.007 | 48.36*** | 0.170 | 0.006 | 29.423*** | 0.320 | 0.007 | 49.223*** |
| Pitch-accented × music lessons | 0.086 | 0.017 | 5.115*** | 0.058 | 0.014 | 4.178*** | -0.007 | 0.016 | -0.46 |
| Tonal × music lessons | -0.020 | 0.013 | -1.508 | -0.012 | 0.011 | -1.095 | 0.067 | 0.012 | 5.486*** |
| Intercept | -0.382 | 0.026 | -14.44*** | -0.008 | 0.025 | -0.31 | -0.219 | 0.018 | -11.889*** |

**Table S5.** Fixed-effect output from random-effects model applied to each musical task controlling for region (East vs West, baseline = East). Language and country were entered as random-effects. Only speakers of tonal and non-tonal languages from countries/regions China, Hong Kong, Taiwan, Thailand, Vietnman (East) and United States, United Kingdom, New Zealand, Canada, Australia (West) are included in this analysis. \*\*\**p* < 0.001, \*\**p* < 0.01, *p < 0.05.

| Term | Melodic Discrimination |  |  | Mistuning Perception |  |  | Beat Alignment |  |  |
| --- | --- | --- | --- | --- | --- | --- | --- | --- | --- |
| | $\beta$ | $SE$ | $t$ | $\beta$ | $SE$ | $t$ | $\beta$ | $SE$ | $t$ |
| **Language: Tonal** | **0.285** | **0.060** | **4.765\*\*\*** | **-0.066** | **0.047** | **-1.403** | **-0.201** | **0.035** | **-5.826\*\*\*** |
| Music lessons: Yes | 0.573 | 0.006 | 96.863*** | 0.441 | 0.005 | 94.699*** | 0.276 | 0.005 | 52.023*** |
| Age | 0.003 | 0.000 | 13.536*** | 0.002 | 0.000 | 12.282*** | -0.003 | 0.000 | -14.772*** |
| Gender: Male | 0.105 | 0.005 | 23.364*** | -0.050 | 0.004 | -14.004*** | 0.128 | 0.004 | 31.699*** |
| Gender: Other | 0.048 | 0.015 | 3.132** | -0.086 | 0.012 | -7.077*** | 0.056 | 0.014 | 4.029*** |
| Region: West | -0.040 | 0.040 | -1.008 | 0.082 | 0.034 | 2.378* | 0.058 | 0.021 | 2.792 |
| Tonal × music lessons | -0.109 | 0.015 | -7.478*** | -0.044 | 0.011 | -3.887*** | 0.052 | 0.013 | 4.038*** |
| Intercept | -0.138 | 0.044 | -3.171** | 0.051 | 0.036 | 1.419 | -0.015 | 0.025 | -0.59 |

**Table S6A.** Summary of main results from linear regressions in the confirmatory sample, repeated across the four analysis approaches (*p < 0.05, ***p < 0.001).

| Term | Melodic Discrimination | Mistuning Perception | Beat Alignment |
| --- | --- | --- | --- |
| **Main Analysis** |  |  |  |
| Language: Pitch-accented | 0.071*** | 0.141*** | 0.173*** |
| Language: Tonal | 0.418*** | -0.094*** | -0.102*** |
| **Matched** |  |  |  |
| Language: Pitch-accented | 0.061*** | 0.128*** | 0.156*** |
| Language: Tonal | 0.351*** | -0.138*** | -0.174*** |
| **Matched (No Lessons Only)** |  |  |  |
| Language: Pitch-accented | 0.044 | 0.174*** | 0.209*** |
| Language: Tonal | 0.437*** | -0.024 | -0.129*** |
| **Inverse Probability Weighted** |  |  |  |
| Language: Pitch-accented | 0.054*** | 0.122*** | 0.154*** |
| Language: Tonal | 0.344*** | -0.136*** | -0.161*** |

**Table S6B.** Beta coefficients from linear regression models for different analysis approaches in the exploratory sample (*p < 0.05, ***p < 0.001).

| Term | Melodic Discrimination | Mistuning Perception | Beat Alignment |
| --- | --- | --- | --- |
| **Main Analysis** |  |  |  |
| Language: Pitch-accented | 0.086*** | 0.158*** | 0.124*** |
| Language: Tonal | 0.403*** | -0.111*** | -0.158*** |
| **Matched** |  |  |  |
| Language: Pitch-accented | 0.074*** | 0.127*** | 0.133*** |
| Language: Tonal | 0.348*** | -0.175*** | -0.185*** |
| **Matched (No Lessons Only)** |  |  |  |
| Language: Pitch-accented | 0.086* | 0.102*** | 0.176*** |
| Language: Tonal | 0.474*** | -0.113*** | -0.188*** |
| **Inverse Probability Weighted** |  |  |  |
| Language: Pitch-accented | 0.082*** | 0.149*** | 0.116*** |
| Language: Tonal | 0.347*** | -0.144*** | -0.209*** |

**Figure S1.**
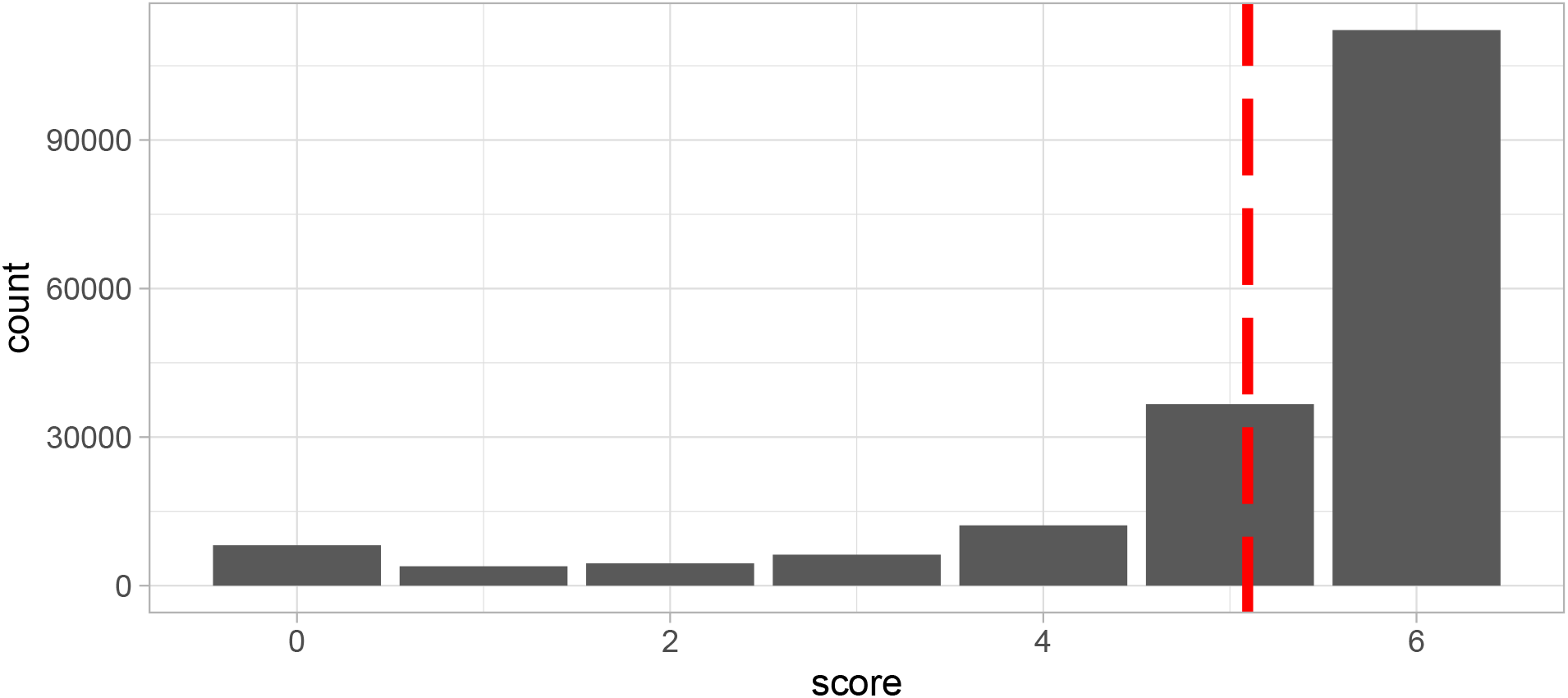
Scores on the headphone detection task, from participants who self-reported that they were wearing headphones. The maximum score was 6; the dashed red line indicates the mean score.

**Figure S2.**
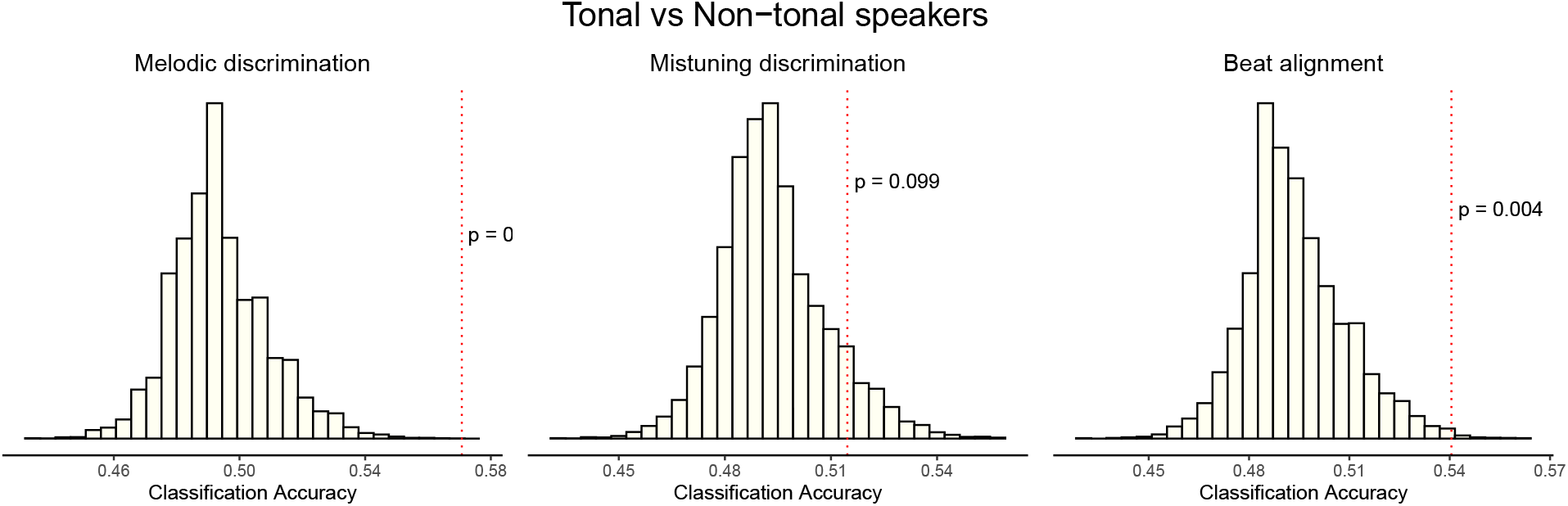
To test whether speakers of tonal and non-tonal languages could be distinguished on the basis of their music perception scores, we ran a permuted Discriminant Function Analysis (pDFA). The three histograms show the null distributions for each music perception task, representing the ability to discriminate ‘tonal’ from ‘non-tonal’ when these labels have been randomly shuffled, resulting from 10,000 permutations. The dotted vertical line then represents the actual discrimination performance on non-shuffled data. And the p-value is computed as a approximate test from this null distribution and actual score.

**Figure S3.**
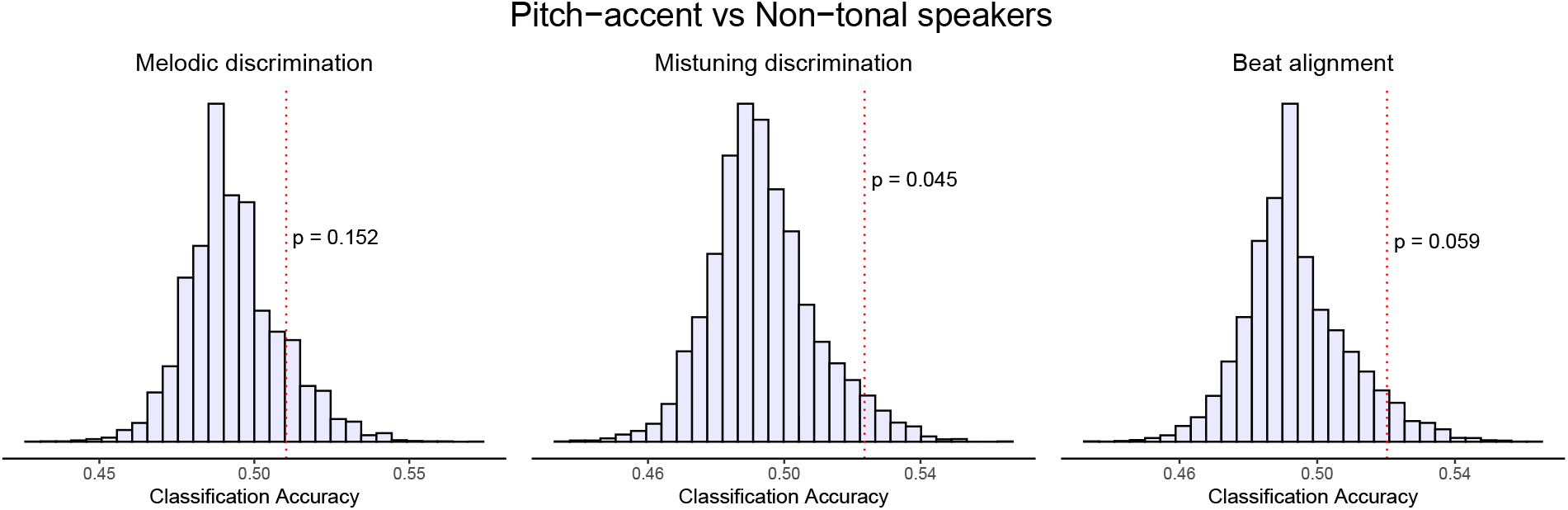
To test whether speakers of pitch-accented and non-tonal languages could be distinguished on the basis of their music perception scores, we ran a permuted Discriminant Function Analysis (pDFA). The three histograms show the null distributions for each music perception task, representing the ability to discriminate ‘tonal’ from ‘non-tonal’ when these labels have been randomly shuffled, resulting from 10,000 permutations. The dotted vertical line then represents the actual discrimination performance on non-shuffled data. And the p-value is computed as a approximate test from this null distribution and actual score.

## Footnotes

1 Indeed, a category of languages that groups evidently disparate languages together, such as Japanese and Swedish, may not be defensible. As pitch-accented languages are not our primary focus, here we treat them as a separate group from tonal and non-tonal languages, but also conduct some analyses at the language level rather than the language-type level.

2 For Jasmin et al. (2021), which studied several participants with lengthy exposure to musical training, we excluded those with more than three years of musical training and recalculated the relevant statistics

## References

Albouy, P., Benjamin, L., Morillon, B., & Zatorre, R. J. (2020). Distinct sensitivity to spectrotemporal modulation supports brain asymmetry for speech and melody. Science, 367 (6481), 1043–1047. https://doi.org/10.1126/science.aaz3468

Alexander, J. A., Bradlow, A. R., Ashley, R. D., & Wong, P. C. M. (2008). Music melody perception in tone-language- and nontone-language speakers. The Journal of the Acoustical Society of America, 124(4), 2495–2495. https://doi.org/10.1121/1.4782815

Ameka, F. K. (2001). Ideophones and the nature of the adjective word class in Ewe. Typological Studies in Language, 44(25-48).

Asano, R., Boeckx, C., & Seifert, U. (2021). Hierarchical control as a shared neurocognitive mechanism for language and music. Cognition. https://doi.org/10.31234/osf.io/25qha

Asaridou, S. S., & McQueen, J. M. (2013). Speech and music shape the listening brain: Evidence for shared domain-general mechanisms. Frontiers in Psychology, 4. https://doi.org/10.3389/fpsyg.2013.00321

Austin, P. C., & Stuart, E. A. (2015). Moving towards best practice when using inverse probability of treatment weighting (IPTW) using the propensity score to estimate causal treatment effects in observational studies. Statistics in Medicine, 34(28), 3661–3679. https://doi.org/10.1002/sim.6607

Bailey, L. M. (1988). A non-linear analysis of pitch accent in Swedish. Lingua, 75(2), 103–124. https://doi.org/10.1016/0024-3841(88)90028-9

Bates, D., Mächler, M., Bolker, B., & Walker, S. (2015). Fitting Linear Mixed-Effects Models Using **Lme4**. Journal of Statistical Software, 67 (1). https://doi.org/10.18637/jss.v067.i01

Bent, T., Bradlow, A. R., & Wright, B. A. (2006). The influence of linguistic experience on the cognitive processing of pitch in speech and nonspeech sounds. Journal of Experimental Psychology: Human Perception and Performance, 32(1), 97–103. https://doi.org/10.1037/0096-1523.32.1.97

Bergelson, E., Amatuni, A., Dailey, S., Koorathota, S., & Tor, S. (2019). Day by day, hour by hour: Naturalistic language input to infants. Developmental Science, 22(1), e12715. https://doi.org/10.1111/desc.12715

Bidelman, G. M., Gandour, J. T., & Krishnan, A. (2011a). Musicians and tone-language speakers share enhanced brainstem encoding but not perceptual benefits for musical pitch. Brain and Cognition, 77 (1), 1–10. https://doi.org/10.1016/j.bandc.2011.07.006

Bidelman, G. M., Gandour, J. T., & Krishnan, A. (2011b). Cross-domain effects of music and language experience on the representation of pitch in the human auditory brainstem. Journal of Cognitive Neuroscience, 23(2), 425–434.

Bidelman, G. M., Hutka, S., & Moreno, S. (2013). Tone Language Speakers and Musicians Share Enhanced Perceptual and Cognitive Abilities for Musical Pitch: Evidence for Bidirectionality between the Domains of Language and Music. PLOS ONE, 8(4), e60676. https://doi.org/10.1371/journal.pone.0060676

Bidelman, G. M., & Lee, C.-C. (2015). Effects of language experience and stimulus context on the neural organization and categorical perception of speech. NeuroImage, 120, 191–200. https://doi.org/10.1016/j.neuroimage.2015.06.087

Blasi, D. E., Henrich, J., Adamou, E., Kemmerer, D., & Majid, A. (2022). Over-reliance on English hinders cognitive science. Trends in Cognitive Sciences.

Bonneville-Roussy, A., Rentfrow, P. J., Xu, M. K., & Potter, J. (2013). Music through the ages: Trends in musical engagement and preferences from adolescence through middle adulthood. Journal of Personality and Social Psychology, 105(4), 703–717. https://doi.org/10.1037/a0033770

Borenstein, M., Hedges, L. V., Higgins, J. P. T., & Rothstein, H. R. (2009). Introduction to meta-analysis. John Wiley & Sons.

Bradley, E. D. (2016). Phonetic dimensions of tone language effects on musical melody perception. Psychomusicology: Music, Mind, and Brain, 26(4), 337–345. https://doi.org/10.1037/pmu0000162

Breen, M., Fedorenko, E., Wagner, M., & Gibson, E. (2010). Acoustic correlates of information structure. Language and Cognitive Processes, 25(7-9), 1044–1098. https://doi.org/10.1080/01690965.2010.504378

Campbell, P. S., & Wiggins, T. (2012). The Oxford Handbook of Children’s Musical Cultures. Oxford University Press.

Chang, D., Hedberg, N., & Wang, Y. (2016). Effects of musical and linguistic experience on categorization of lexical and melodic tones. The Journal of the Acoustical Society of America, 139(5), 2432–2447. https://doi.org/10.1121/1.4947497

Chen, A., Liu, L., & Kager, R. (2016). Cross-domain correlation in pitch perception, the influence of native language. Language, Cognition and Neuroscience, 31(6), 751–760. https://doi.org/10.1080/23273798.2016.1156715

Choi, W. (2021). Musicianship Influences Language Effect on Musical Pitch Perception. Frontiers in Psychology, 12, 712753. https://doi.org/10.3389/fpsyg.2021.712753

Cowen, A. S., Laukka, P., Elfenbein, H. A., Liu, R., & Keltner, D. (2019). The primacy of categories in the recognition of 12 emotions in speech prosody across two cultures. Nature Human Behaviour, 3(4), 369–382. https://doi.org/10.1038/s41562-019-0533-6

de Leeuw, J. R. (2015). jsPsych: A JavaScript library for creating behavioral experiments in a Web browser. Behavior Research Methods, 47 (1), 1–12. https://doi.org/10.3758/s13428-014-0458-y

Delogu, F., Lampis, G., & Belardinelli, M. O. (2010). From melody to lexical tone: Musical ability enhances specific aspects of foreign language perception. European Journal of Cognitive Psychology, 22(1), 46–61. https://doi.org/10.1080/09541440802708136

Delogu, F., Lampis, G., & Olivetti Belardinelli, M. (2006). Music-to-language transfer effect: May melodic ability improve learning of tonal languages by native nontonal speakers? Cognitive Processing, 7 (3), 203–207. https://doi.org/10.1007/s10339-006-0146-7

Eibl-Eibesfeldt, I. (1979). Human ethology: Concepts and implications for the sciences of man. Behavioral and Brain Sciences, 2(01), 1–57. https://doi.org/10.1017/S0140525X00060416

Eme, C. A., & Odinye, I. S. (2022). Phonology of standard Chinese and Igbo: Implications for Igbo students learning Chinese. PUBLISHED ACADEMIC PAPERS, 0.

Evans, J., Yeh, W.-C., & Kulkarni, R. (2018). Acoustics of Tone in Indian Punjabi. Transactions of the Philological Society, 116(3), 509–528. https://doi.org/10.1111/1467-968X.12135

Giuliano, R. J., Pfordresher, P. Q., Stanley, E. M., Narayana, S., & Wicha, N. Y. Y. (2011). Native Experience with a Tone Language Enhances Pitch Discrimination and the Timing of Neural Responses to Pitch Change. Frontiers in Psychology, 2. https://doi.org/10.3389/fpsyg.2011.00146

Goldsmith, J., & Mpiranya, F. (2011). Rhythm, quantity and tone in the Kinyarwanda verb. In J. A. Goldsmith, E. Hume, & L. Wetzels (Eds.), Tones and Features (pp. 25–49). DE GRUYTER. https://doi.org/10.1515/9783110246223.25

Gussenhoven, C. (2004). The Phonology of Tone and Intonation. Cambridge University Press.

Hamann, S., & Kula, N. C. (2015). Bemba. Journal of the International Phonetic Association, 45(1), 61–69. https://doi.org/10.1017/S0025100314000371

Hannon, E. E., & Trehub, S. E. (2005). Metrical categories in infancy and adulthood. Psychological Science, 16(1), 48–55.

Harrison, P. M. c. (2020). psychTestR: An R package for designing and conducting behavioural psychological experiments. Journal of Open Source Software, 5(49), 2088. https://doi.org/10.21105/joss.02088

Harrison, P. M. C., Collins, T., & Müllensiefen, D. (2017). Applying modern psychometric techniques to melodic discrimination testing: Item response theory, computerised adaptive testing, and automatic item generation. Scientific Reports, 7 (1), 1–18. https://doi.org/10.1038/s41598-017-03586-z

Harrison, P. M. C., & Müllensiefen, D. (2018). Development and validation of the Computerised Adaptive Beat Alignment Test (CA-BAT). Scientific Reports, 8(1), 1–19. https://doi.org/10.1038/s41598-018-30318-8

Hartshorne, J. K., de Leeuw, J., Goodman, N., Jennings, M., & O’Donnell, T. J. (2019). A thousand studies for the price of one: Accelerating psychological science with Pushkin. Behavior Research Methods, 51, 1782–1803. https://doi.org/10.3758/s13428-018-1155-z

Hilton, C. B., & Goldwater, M. (2021). Linguistic syncopation: Meter-syntax alignment affects sentence comprehension and sensorimotor synchronization. Cognition. https://doi.org/10.31234/osf.io/hcngm

Hilton, C. B., & Mehr, S. A. (2022). Citizen science can help to alleviate the generalizability crisis. Behavioral and Brain Sciences.

Hilton, C. B., Moser, C. J., Bertolo, M., Lee-Rubin, H., Amir, D., Bainbridge, C. M., Simson, J., Knox, D., Glowacki, L., Alemu, E., Galbarczyk, A., Jasienska, G., Ross, C. T., Neff, M. B., Martin, A., Cirelli, L. K., Trehub, S. E., Song, J., Kim, M., … Mehr, S. A. (2022). Acoustic regularities in infant-directed speech and song across cultures. Nature Human Behaviour. https://doi.org/10.1101/2020.04.09.032995

Huber, B., & Gajos, K. Z. (2020). Conducting online virtual environment experiments with uncompensated, unsupervised samples. PLOS ONE, 15(1), e0227629. https://doi.org/10.1371/journal.pone.0227629

Hutka, S., Bidelman, G. M., & Moreno, S. (2015). Pitch expertise is not created equal: Cross-domain effects of musicianship and tone language experience on neural and behavioural discrimination of speech and music. Neuropsychologia, 71, 52–63. https://doi.org/10.1016/j.neuropsychologia.2015.03.019

Hyman, L. M. (2009). How (not) to do phonological typology: The case of pitch-accent. Language Sciences, 31(2), 213–238. https://doi.org/10.1016/j.langsci.2008.12.007

Hyman, L. M. (2006). Word-Prosodic Typology. Phonology, 23(2), 225–257.

Inkelas, S., & Zec, D. (1988). Serbo-Croatian Pitch Accent: The Interaction of Tone, Stress, and Intonation. Language, 64(2), 227–248. https://doi.org/10.2307/415433

Jasmin, K., Dick, F., Stewart, L., & Tierney, A. T. (2020). Altered functional connectivity during speech perception in congenital amusia. eLife, 9, e53539. https://doi.org/10.7554/eLife.53539

Jasmin, K., Sun, H., & Tierney, A. T. (2020). Effects of language experience on domain-general perceptual strategies. bioRxiv, 2020.01.02.892943. https://doi.org/10.1101/2020.01.02.892943

Jasmin, K., Sun, H., & Tierney, A. T. (2021). Effects of language experience on domain-general perceptual strategies. Cognition, 206, 104481. https://doi.org/10.1016/j.cognition.2020.104481

Jefferies, A. A. (1990). Beyond tone: Functions of pitch in Shona [PhD thesis]. University of Florida.

Konner, M. (2010). The evolution of childhood: Relationships, emotion, mind. Belknap Press of Harvard University Press.

Kraemer, H. C., Gardner, C., Brooks III, J. O., & Yesavage, J. A. (1998). Advantages of excluding underpowered studies in meta-analysis: Inclusionist versus exclusionist viewpoints. Psychological Methods, 3(1), 23. https://doi.org/10.1037/1082-989X.3.1.23

Kraus, N., & Chandrasekaran, B. (2010). Music training for the development of auditory skills. Nature Reviews Neuroscience, 11(8), 599–605.

Krishnan, A., Gandour, J. T., & Bidelman, G. M. (2010). The effects of tone language experience on pitch processing in the brainstem. Journal of Neurolinguistics, 23(1), 81–95. https://doi.org/10.1016/j.jneuroling.2009.09.001

Krishnan, A., Xu, Y., Gandour, J., & Cariani, P. (2005). Encoding of pitch in the human brainstem is sensitive to language experience. Cognitive Brain Research, 25(1), 161–168. https://doi.org/10.1016/j.cogbrainres.2005.05.004

Krizman, J., Marian, V., Shook, A., Skoe, E., & Kraus, N. (2012). Subcortical encoding of sound is enhanced in bilinguals and relates to executive function advantages. Proceedings of the National Academy of Sciences, 109(20), 7877–7881. https://doi.org/10.1073/pnas.1201575109

Krumhansl, C. L. (2004). The Cognition of Tonality – as We Know it Today. Journal of New Music Research, 33(3), 253–268. https://doi.org/10.1080/0929821042000317831

Kuhl, P. K. (2004). Early language acquisition: Cracking the speech code. Nature Reviews Neuroscience, 5(11), 831–843. https://doi.org/10.1038/nrn1533

Larrouy-Maestri, P., Harrison, P. M. C., & Müllensiefen, D. (2019). The mistuning perception test: A new measurement instrument. Behavior Research Methods, 51(2), 663–675. https://doi.org/10.3758/s13428-019-01225-1

Li, X., & Gao, Z. (2018). The effect of language experience on lexical tone perception. The Journal of the Acoustical Society of America, 144(3), 1865–1865. https://doi.org/10.1121/1.5068200

Liu, L., Chen, A., & Kager, R. (2020). Simultaneous bilinguals who do not speak a tone language show enhancement in pitch sensitivity but not in executive function. https://doi.org/10.1075/lab.19037.liu

Liu, L., & Kager, R. (2017). Enhanced music sensitivity in 9-month-old bilingual infants. Cognitive Processing, 18(1), 55–65. https://doi.org/10.1007/s10339-016-0780-7

Long, B., Simson, J., Buxó-Lugo, A., Watson, D. G., & Mehr, S. A. (2023). How games can make behavioural science better. Nature, 613(7944), 433–436. https://doi.org/10.1038/d41586-023-00065-6

Lynch, M. P., Eilers, R. E., Oller, D. K., & Urbano, R. C. (1990). Innateness, Experience, and Music Perception. Psychological Science, 1(4), 272–276. https://doi.org/10.1111/j.1467-9280.1990.tb00213.x

Maddieson, I. (2013). Tone. In The World Altas of Language Structures Online (Dryer, Matthew S. & Haspelmath, Martin).

Maddieson, I., Flavier, S., & Pellegrino, F. (2014). LAPSyD: Lyon-Albuquerque phonological systems databases, Version 1.0.

Manyah, K. A. (2006). Relation between tone and vowel quality in Twi. International Symposium on Tonal Aspects of Languages.

McMullen, E., & Saffran, J. R. (2004). Music and Language: A Developmental Comparison. Music Perception, 21(3), 289–311. https://doi.org/10.1525/mp.2004.21.3.289

Mehr, S. A. (2014). Music in the home: New evidence for an intergenerational link. Journal of Research in Music Education, 62(1), 78–88. https://doi.org/10.1177/0022429413520008

Mehr, S. A., Krasnow, M. M., Bryant, G. A., & Hagen, E. H. (2020). Origins of music in credible signaling. Behavioral and Brain Sciences, 1–41. https://doi.org/10.1017/S0140525X20000345

Mehr, S. A., Schachner, A., Katz, R. C., & Spelke, E. S. (2013). Two randomized trials provide no consistent evidence for nonmusical cognitive benefits of brief preschool music enrichment. PLoS ONE, 8(12), e82007. https://doi.org/10.1371/journal.pone.0082007

Mehr, S. A., Singh, M., Knox, D., Ketter, D. M., Pickens-Jones, D., Atwood, S., Lucas, C., Jacoby, N., Egner, A. A., Hopkins, E. J., Howard, R. M., Hartshorne, J. K., Jennings, M. V., Simson, J., Bainbridge, C. M., Pinker, S., O’Donnell, T. J., Krasnow, M. M., & Glowacki, L. (2019). Universality and diversity in human song. Science, 366(6468), 957–970. https://doi.org/10.1126/science.aax0868

Mendoza, J. K., & Fausey, C. M. (2021). Everyday music in infancy. Developmental Science. https://doi.org/10.31234/osf.io/sqatb

Moran, S., & McCloy, D. (Eds.). (2019). PHOIBLE 2.0. Max Planck Institute for the Science of Human History.

Mundry, R., & Sommer, C. (2007). Discriminant function analysis with nonindependent data: Consequences and an alternative. Animal Behaviour, 74(4), 965–976. https://doi.org/10.1016/j.anbehav.2006.12.028

Ngo, M. K., Vu, K.-P. L., & Strybel, T. Z. (2016). Effects of Music and Tonal Language Experience on Relative Pitch Performance. The American Journal of Psychology, 129(2), 125–134. https://doi.org/10.5406/amerjpsyc.129.2.0125

Niesler, T., Louw, P., & Roux, J. (2005). Phonetic analysis of Afrikaans, English, Xhosa and Zulu using South African speech databases. Southern African Linguistics and Applied Language Studies, 23(4), 459–474. https://doi.org/10.2989/16073610509486401

Patel, A. D. (2012). The OPERA hypothesis: Assumptions and clarifications. Annals of the New York Academy of Sciences, 1252(1), 124–128. https://doi.org/10.1111/j.1749-6632.2011.06426.x

Patel, A. D. (2008). Music, language, and the brain. Oxford University Press.

Patel, A. D. (2011). Why would musical training benefit the neural encoding of speech? The OPERA hypothesis. Frontiers in Psychology, 2.

Peng, G., Zheng, H.-Y., Gong, T., Yang, R.-X., Kong, J.-P., & Wang, W. S.-Y. (2010). The influence of language experience on categorical perception of pitch contours. Journal of Phonetics, 38(4), 616–624. https://doi.org/10.1016/j.wocn.2010.09.003

Peretz, I., Champod, A. S., & Hyde, K. (2003). Varieties of musical disorders. The Montreal Battery of Evaluation of Amusia. Annals of the New York Academy of Sciences, 999(1), 58–75. https://doi.org/10.1196/annals.1284.006

Peretz, I., Nguyen, S., & Cummings, S. (2011). Tone Language Fluency Impairs Pitch Discrimination. Frontiers in Psychology, 2. https://doi.org/10.3389/fpsyg.2011.00145

Peretz, I., Vuvan, D., Lagrois, M.-É., & Armony, J. L. (2015). Neural overlap in processing music and speech. Philosophical Transactions of the Royal Society of London B: Biological Sciences, 370(1664), 20140090. https://doi.org/10.1098/rstb.2014.0090

Pfordresher, P. Q., & Brown, S. (2009). Enhanced production and perception of musical pitch in tone language speakers. Attention, Perception, & Psychophysics, 71(6), 1385–1398. https://doi.org/10.3758/APP.71.6.1385

Pike, K. L. (1948). Tone languages. University of Michigan Press.

Polka, L., & Werker, J. F. (1994). Developmental changes in perception of nonnative vowel contrasts. Journal of Experimental Psychology: Human Perception and Performance, 20(2), 421–435. https://doi.org/10.1037/0096-1523.20.2.421

Sala, G., & Gobet, F. (2020). Cognitive and academic benefits of music training with children: A multilevel meta-analysis. Memory & Cognition, 48(8), 1429–1441. https://doi.org/10.3758/s13421-020-01060-2

Schertz, J., & Clare, E. J. (2020). Phonetic cue weighting in perception and production. WIREs Cognitive Science, 11(2), e1521. https://doi.org/10.1002/wcs.1521

Stevens, C. J., Keller, P. E., & Tyler, M. D. (2013). Tonal language background and detecting pitch contour in spoken and musical items. Psychology of Music, 41(1), 59–74. https://doi.org/10.1177/0305735611415749

Stuart, E. A. (2010). Matching methods for causal inference: A review and a look forward. Statistical Science : A Review Journal of the Institute of Mathematical Statistics, 25(1), 1–21. https://doi.org/10.1214/09-STS313

Swaminathan, S., Kragness, H. E., & Schellenberg, E. G. (2021). The Musical Ear Test: Norms and correlates from a large sample of Canadian undergraduates. Behavior Research Methods. https://doi.org/10.3758/s13428-020-01528-8

Tierney, A., Krizman, J., Skoe, E., Johnston, K., & Kraus, N. (2013). High school music classes enhance the neural processing of speech. Frontiers in Psychology, 4. https://doi.org/10.3389/fpsyg.2013.00855

Tong, X., Choi, W., & Man, Y. Y. (2018). Tone language experience modulates the effect of long-term musical training on musical pitch perception. The Journal of the Acoustical Society of America, 144(2), 690–697. https://doi.org/10.1121/1.5049365

Tong, Y., Gandour, J., Talavage, T., Wong, D., Dzemidzic, M., Xu, Y., Li, X., & Lowe, M. (2005). Neural circuitry underlying sentence-level linguistic prosody. NeuroImage, 28(2), 417–428. https://doi.org/10.1016/j.neuroimage.2005.06.002

van der Hulst, H. (2011). Pitch Accent Systems. In The Blackwell Companion to Phonology (pp. 1–24). Wiley-Blackwell. https://doi.org/10.1002/9781444335262.wbctp0042

van der Hulst, H., Goedemans, R., & van Zanten, E. (2010). A Survey of Word Accentual Patterns in the Languages of the World. De Gruyter Mouton.

Wagner, M., & McAuliffe, M. (2019). The effect of focus prominence on phrasing. Journal of Phonetics, 77, 100930. https://doi.org/10.1016/j.wocn.2019.100930

Wang, Q. (2008). L2 stress perception: The reliance on different acoustic cues. Proceedings of Speech Prosody, 135–138. https://doi.org/https://www.isca-speech.org/archive/sp2008/papers/sp08_635.pdf

Werker, J. F., & Tees, R. C. (1984). Cross-language speech perception: Evidence for perceptual reorganization during the first year of life. Infant Behavior and Development, 7 (1), 49–63. https://doi.org/10.1016/S0163-6383(84)80022-3

Westermeyer, R., & Westermeyer, J. (1977). Tonal Language Acquisition among Lao Children. Anthropological Linguistics, 19(6), 260–264.

Wong, P. C. M., Ciocca, V., Chan, A. H. D., Ha, L. Y. Y., Tan, L.-H., & Peretz, I. (2012). Effects of Culture on Musical Pitch Perception. PLOS ONE, 7 (4), e33424. https://doi.org/10.1371/journal.pone.0033424

Wong, P. C. M., Kang, X., Wong, K. H. Y., So, H.-C., Choy, K. W., & Geng, X. (2020). ASPM-lexical tone association in speakers of a tone language: Direct evidence for the genetic-biasing hypothesis of language evolution. Science Advances, 6(22), eaba5090. https://doi.org/10.1126/sciadv.aba5090

Wong, P. C. M., Skoe, E., Russo, N. M., Dees, T., & Kraus, N. (2007). Musical experience shapes human brainstem encoding of linguistic pitch patterns. Nature Neuroscience, 10(4), 420–422. https://doi.org/10.1038/nn1872

Woods, K. J. P., Siegel, M. H., Traer, J., & McDermott, J. H. (2017). Headphone screening to facilitate web-based auditory experiments. Attention, Perception, & Psychophysics, 1–9. https://doi.org/10.3758/s13414-017-1361-2

Yan, R., Jessani, G., Spelke, E., Villiers, P. de, Villiers, J. de, & Mehr, S. (2021). Across demographics and recent history, most parents sing to their infants and toddlers daily. Philosophical Transactions of the Royal Society B: Biological Sciences. https://doi.org/10.31234/osf.io/fy5bh

Yarkoni, T. (2022). The generalizability crisis. Behavioral and Brain Sciences, 45. https://doi.org/10.1017/S0140525X20001685

Yip, M. (2002). Tone. Cambridge University Press.

Yu, V. Y., & Andruski, J. E. (2010). A Cross-Language Study of Perception of Lexical Stress in English. Journal of Psycholinguistic Research, 39(4), 323–344. https://doi.org/10.1007/s10936-009-9142-2

Zatorre, R. J., Belin, P., & Penhune, V. B. (2002). Structure and function of auditory cortex: Music and speech. Trends in Cognitive Sciences, 6(1), 37–46. https://doi.org/10.1016/S1364-6613(00)01816-7

Zhang, L., Xie, S., Li, Y., Shu, H., & Zhang, Y. (2020). Perception of musical melody and rhythm as influenced by native language experience. The Journal of the Acoustical Society of America, 147 (5), EL385–EL390. https://doi.org/10.1121/10.0001179

Zheng, Y., & Samuel, A. G. (2018). The effects of ethnicity, musicianship, and tone language experience on pitch perception. Quarterly Journal of Experimental Psychology, 71(12), 2627–2642. https://doi.org/10.1177/1747021818757435

